# Integrative multiomic analysis on single-nucleotide variants identifies candidate genes for human craniofacial malformation

**DOI:** 10.64898/2025.12.29.696805

**Authors:** Ming Ho Yam, Karl Kam Hei So, Ka Kui Tong, Kwong Wai Choy, Mai Har Sham

**Author notes:** ¶ These authors contributed equally to this work.

## Abstract

Craniofacial malformation (CFM) is a congenital defect encompassing a wide range of phenotypic presentations and is largely driven by genetics. Despite the discovery of more than 300 causal genes, there are a myriad of CFM cases with unknown genetic etiology. The complex gene regulations and heterogeneous cellular interactions in the developing head complicate disease-gene identification and prenatal genetic diagnosis. Recent progress in multiomic profiling of human embryogenesis enables the discovery of novel candidates from established GWAS data. Here, we developed an approach to prioritize GWAS variants using the epigenomes and single-cell transcriptomes of embryonic tissues and progenitor cells by implementing machine learning classifiers and combinatorial analysis. Systematic evaluation revealed significant improvement in the machine learning model performance after integrating transcriptome of neural crest cells (NCCs) and cranial placodes, as well as epigenomic profile of early craniofacial tissues. We identified 249 genes from the best-performing classifier, which include documented CFM-associated genes. Gene regulatory network (GRN) inference showed that 24 candidate genes were involved in NCC- and placode-specific regulons, of which 15 (*F11R, ISL1, KANK4, L1TD1, LAMB1, MIA, PRDM1, S100A10, S100A11, STOM, STT3B, TESK2, USP43, WDR86, ZNF439*) were novel candidates for human CFM. Motif analysis revealed putative functional SNPs contributing to CFM pathogenesis by disrupting transcription factor binding motifs in neural crest and placodes. Our analyses suggested that *PRDM1* and *ISL1* are strong candidates for human CFM, as supported by other animal functional studies. This study demonstrates a successful method for disease gene identification using epigenomic and single-cell transcriptomic profiles, and sheds light on the linkage between early cell lineages and the pathogenic process of CFM.

**Author Summary:** Craniofacial malformation is one of the most common congenital disorders that affects food ingestion, speech and social interaction of the patients. The identification of craniofacial disease genes is difficult due to the dynamic gene expression and contribution from multiple cell types during embryonic development. In this study, we combine artificial intelligence with patient genetic and embryo multiomic information to identify new candidate genes for human craniofacial malformation. Using machine learning classifiers and combinatorial analyses, we prioritized single-nucleotide variants from patient datasets and identified 249 candidate genes. Annotation of the variants and candidate genes showed that some of them overlapped with known disease genes, demonstrating the efficacy of our approach. Further analyses using lineage reconstruction and motif analyses revealed a number of promising novel candidates, in particular *PRDM1* and *ISL1*, are likely to be causative genes for human CFM. Our study has demonstrated a translatable approach for disease gene identification utilizing machine learning algorithm and multiomic data, and provides a gene list for improving diagnostic panels and understanding the pathogenic processes of craniofacial disorders.

## Introduction

Craniofacial malformation (CFM) is one of the most common congenital disorders affecting facial structures and functions. Patients with CFM exhibit a broad spectrum of clinical presentations, including orofacial cleft, micrognathia, and craniosynostosis (1). Both normal and malformed facial variations are largely determined by genetic factors (2). Understanding the genetic basis of CFM is essential not only for understanding the pathogenic processes, but also for improving clinical diagnosis and classification of syndromic disorders. Nevertheless, in many clinical cases, the underlying genetic causes remain unknown. This is partly due to the polygenic nature of CFM and the phenotypic heterogeneity presented by patients, complicating the establishment of genotype-phenotype correlations.

Over the past decades, a number of genomic loci associated with craniofacial variation and diseases have been identified by genome-wide association studies (GWAS) (3–7). Most disease-associated variants reside in non-protein-coding regions, among which a functional subset affects gene expression by disrupting transcription-factor binding to tissue-specific cis-regulatory elements (8). Aberrant activity of these cis-regulatory elements can compromise developmental processes and contribute to morphological defects. A Well-described example is the Pierre Robin sequence (PRS), a craniofacial disorder characterized by anomalies in the mandible and palate (9). PRS-associated mutations in the distal enhancers of *SOX9* are known to reduce *SOX9* expression and impair mandibular morphogenesis (10). Another example is the non-syndromic cleft lips with/without cleft palate. A multi-species comparative study has shown that a common variant (rs2235371) compromised the function of an *IRF6* enhancer by disrupting a *TFAP2A* transcription factor binding site (11). As *TFAP2A* and *IRF6* are connected in the same regulatory cascade underlying the pathogenesis of non-syndromic cleft lips with/without cleft palate, both genes can be considered CFM genes. Multiomic studies further revealed that CFM disease variants are enriched in active regulatory regions in proximity to genes expressed in developing craniofacial tissues (12,13). Importantly, in congenital defects, enhancer-gene interactions take place with temporal and cell-type specificities during fetal development, highlighting the need for integrating epigenomic and single-cell transcriptomic data of embryonic and fetal tissues to identify regulatory gene networks for non-coding disease variants.

The first major event of facial morphogenesis occurs at the fourth week of gestation, marked by the formation of the first pharyngeal arch and frontonasal prominence of the developing head (2,14). The cell types that contribute to these structures are formed during early embryogenesis, which include the multipotent cranial neural crest cells (NCCs) and cranial placodes. Both cranial NCCs and placodes originate from the neural plate border (NPB) between the neural plate and non-neural ectoderm. Cranial NCCs undergo epithelial-to-mesenchymal transition, delaminate from the dorsal neural tube, and migrate to populate the facial prominences. These NCCs contribute to the majority of the craniofacial components by differentiating into chondrocytes and connective tissues. Disrupted neural crest functions cause CFM such as Treacher Collins syndrome, Pierre Robin sequence, and waardenburg syndrome. On the other hand, cranial placodes emerge from the cranial region of the NPB and maintain as epithelial cells. The pre-placodal progenitors proliferate and segregate into distinct epithelial patches, forming transient signaling epithelia and facial sensory organs (15,16). Although less well-recognized, placodal cells are implicated in CFM. Mutations in *SIX1* and *EYA1*, the key regulators of placode development, can cause branchio-oto syndrome (Online Mendelian Inheritance in Man (OMIM): 608389) and oto-facio-cervical syndrome (OMIM: 166780), both exhibiting craniofacial malformation phenotypes. However, due to the scarcity of human samples and the lack of time-resolved epigenomic and single-cell transcriptomic information, the contributions of these early cell lineages have not been extensively investigated in disease gene identification studies. Recently, several transcriptomic and epigenomic atlases of human craniofacial tissues have been established, enabling us to analyse GWAS data With embryonic functional genomic information (12,13,17–20). Advances in single-cell RNA sequencing (scRNA-seq) and software tools also facilitate the identification and investigation of transient cellular populations, including neural crest and placodal cell lineages, during craniofacial development (21,22).

In this study, we integrated epigenomic profiles of human embryos and single-cell transcriptomes of early cell lineages with patient SNP array data to prioritize candidate disease genes. Using machine learning classifiers, we systematically evaluated the disease prediction power of multiple SNP sets, each prioritized by a distinct combination of functional genomic data. This approach identified 249 candidate genes from the best-performing classifier for further analysis, providing a foundation for improving diagnostic gene panels and understanding the pathogenesis of CFM. Our strategy demonstrates the effectiveness of leveraging transcriptomic information from developmentally relevant cell types for disease gene prioritization.

## Result

### Integrating SNP array and functional genomics improves disease classification

We collected SNP arrays from patients with craniofacial disorders and integrated them with functional genomic data from human embryos, in order to prioritize disease-associated variants (Fig 1A). The patient cohort used in this study was diagnosed with craniofacial microsomia, clinically primarily characterized by mandibular hypoplasia and microtia, with additional complications in some cases, including temporal bone agenesis, orofacial clefts, and Treacher Collins syndrome (6). For functional genomic data, to identify regulatory regions associated with actively expressed genes at early time points, we obtained scRNA-seq data from post-conceptual week (PCW) 3-4 whole embryos (20). Ectodermal derivatives were isolated from the processed scRNA-seq data using cellular markers and underwent lineage reconstruction and Leiden clustering (Fig 1B and S1A-C Fig). The ectodermal clusters were annotated into 11 populations based on marker gene expression (S1D Fig). Cell clusters along the NCC and placode lineage trajectories including NCC, placode, non-neural ectoderm and epidermal cells were extracted for disease gene prioritization analysis by retrieving differentially expressed genes (DEGs). In addition, histone modification chromatin immunoprecipitation sequencing (ChIP-Seq) data from PCW 4-5 human craniofacial tissues were processed to identify spatiotemporally specific cis-regulatory elements (Fig 1C) (12). In total, we included ten layers of functional genomics information, comprising six histone modification marks and four sets of DEGs from single-cell transcriptome.

**Fig 1.**
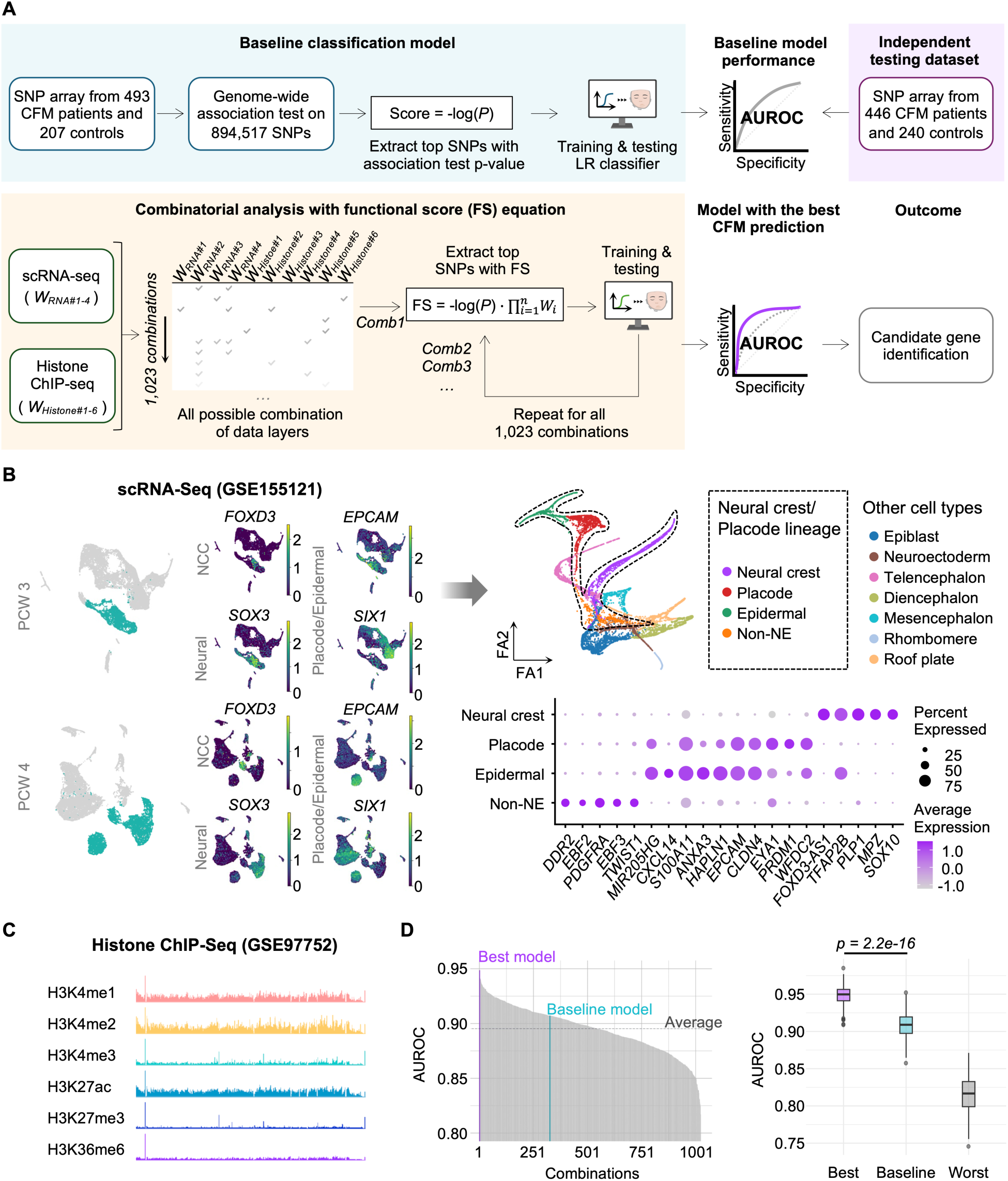
Construction of an integrative multiomic pipeline for SNP prioritization. (A) Overview of data integration workflow. The region in blue demonstrates the process of training the baseline classifier, which is tested with an independent dataset (purple) and assessed using the area under receiver operating characteristic (AUROC) curve. The workflow for combinatorial analysis and functional score (FS) calculated is visualized in the yellow region. The model with the highest AUROC from the combinatorial analysis is used for disease gene identification. (B) Processing of scRNA-Seq data. The left panel displayed the Leiden clustering of scRNA-seq post-conceptual week (PCW) 3-4 embryos and the expression of neural crest (*FOXD3*), neural (*SOX3*), and placode/epidermal (*SIX1*/*EPCAM*) cell markers. The ectodermal subsets are highlighted in green. The Force Atlas plot on the right shows the lineage trajectory of the ectodermal subset with 11 annotated cell populations. Four cell types along the neural crest/placode trajectories (Neural crest, placode, epidermal, and non-neuroepithelium/non-NE) were highlighted and the expression of their top markers was shown in the dot plot. (C) Genome browser tracks showing the histone ChIP-Seq data included for variant prioritization. (D) Mean AUROC of baseline and 1,023 data-integrated classifiers. The best-performing, baseline, and worst-performing models were shown in boxplot.

Candidate SNPs were prioritized following the workflow in Figure 1A. we designed the functional score (FS), a weighted formula that multiplies genome-wide association test p-values by weights from functional genomic annotations. Given that chromatin states and gene expression patterns vary dynamically across cell types during development, we expected that not all data layers would contribute equally to disease gene identification. To systematically identify the most informative combination, we implemented the logistic regression classifier, a type of supervised learning algorithm, to predict disease status using the alleles of prioritized SNPs. The prediction performance was quantified with the average area under receiver operating characteristic curves (AUROC) in 200 iterations of random train-test splits. The model allowed us to evaluate which SNP sets, prioritized by different functional genomic data combinations, were most relevant to the disease. we reasoned that a more biologically informed set of SNPs would result in better disease classification accuracy.

We constructed 1,023 logistic regression classifiers using all possible combinations of functional genomic data, along with a baseline model that did not integrate any additional information. The baseline model, which relied solely on SNP array data, achieved an average AUROC of 0.907 (0.858-0.952, SD = 1.75e-2). Models with additional functional genomics data produced average AUROCs ranging from 0.817 to 0.948, with only 325 models (about one-third) surpassing the baseline performance (Fig 1D). The SNP set in the best-performing classifier was prioritized using NCC and placode DEGs in combination with three histone modification marks (H3K4me3, H3K27me3, H3K36me3). This model exhibited an average AUROC of 0.949 (0.909-0.986, SD = 1.33e-2), significantly higher than the baseline (*p*-value = 2.2e-16) (Fig 1E). Among the top 20-performing combinations, H3K36me3, neural crest DEGs and placode DEGs are the most frequently included data layers (S2 Fig). To evaluate whether integrating functional genomics data preferentially prioritized variants with gene regulatory potential, we utilized two databases, RegulomeDB and FORGEdb, to quantify the functional significance of our variants (23,24). The SNPs that constitute the best-performing classifier exhibit significantly higher RegulomeDB ranks and FORGEdb scores compared with the baseline model, indicating enrichment for regulatory variants (S3 Fig).

### Prioritized SNPs from the best-performing classifier

We retrieved the 137 prioritized SNPs from the classifier with the highest AUROC. They were mapped onto 54 loci in the hg19 genome build (Fig 2 and Table 1). Nine loci contained more than three prioritized SNPs, including 1q21.3, 1q22, 1q23.3, 7q22.1, 7q31.1, 10p14, 17p13.1, 19p13.2, and 19q13.11. To determine whether these candidate loci overlapped with previously reported regions, we identified known disease-associated loci from GWAS Catalog for the phenotypes documented in our SNP array patient cohort, including craniofacial microsomia, outer ear morphological trait, and cleft lip with/without cleft palate (25). Thirty-two out of fifty-four (60%) of our candidate loci were associated with these craniofacial conditions. Notably, the regions 2q37.2 and 10p14 have been reported as genome-wide significant loci for craniofacial microsomia, which is the major feature of our patient cohort (6). The colocalization of prioritized SNPs with known craniofacial disorder genomic regions indicates that our integrative machine learning approach can identify disease-relevant candidates.

**Fig 2.**
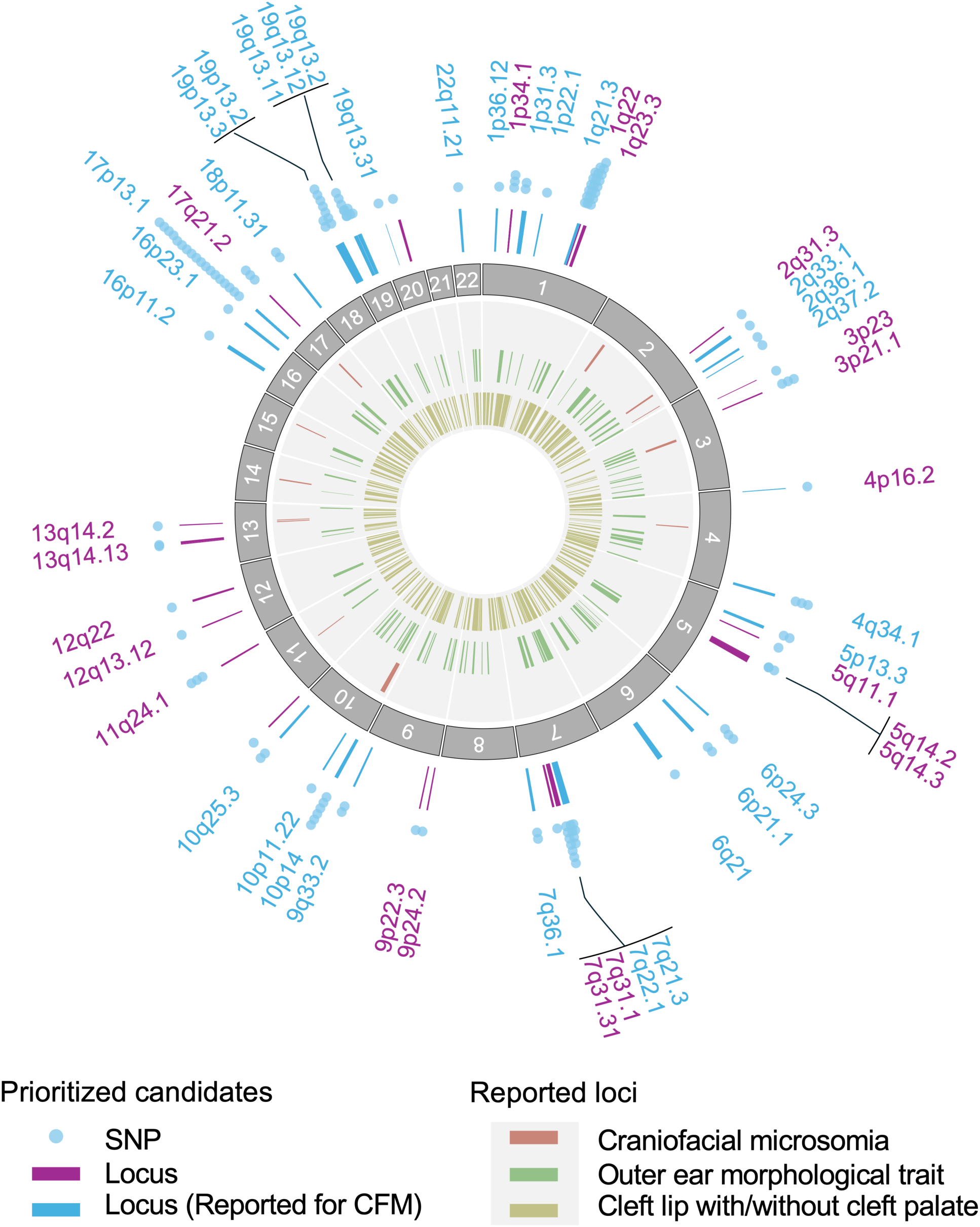
Candidate variants prioritized by the best combination of functional genomic data. Circos plot visualizing prioritized candidates and known associated loci for CFM. The outer region displays the prioritized SNPs and their residing loci. The 137 prioritized SNPs were shown by blue dots. The purple and blue arcs visualize the corresponding loci of each 137 SNPs. The blue arcs indicate that the candidate loci are CFM-associated as documented by human databases. The inner grey region displays known associated loci for craniofacial microsomia (Experimental Factor Ontology (EFO) ID: MONDO_0015397), outer ear morphological trait (EFO ID: EFO_0007664) and cleft lip with/without cleft palate (EFO ID: EFO_0003959).

**Table 1.** The list of SNPs and their corresponding loci from the best-performing machine learning classifier. The first column displays the 54 overlapping loci after mapping the 137 prioritized SNPs to the hg19 genome. The number of SNPs overlapping with each locus is shown in the second column. The rsID of SNPs and their nearest genes are listed in corresponding order in the third and fourth columns respectively.

| Locus | SNP Count | Prioritized SNP (rsID) | Nearest Gene |
| --- | --- | --- | --- |
| 17p13.1 | 18 | rs13342232, rs11656323, rs222843, rs2074217, rs116903547, rs35776863, rs2269762, rs17882626, rs8069501, rs6608, rs11078697, rs28625968, rs9901673, rs858520, rs6503227, rs34654539, rs9896200, rs16958920 | *SLC16A11*, *GABARAP* (x3), *ENSG00000262302*, *NEURL4*, *TNK1*, *TMEM256*, *POLR2A*, *SENP3* (x2), *EIF4A1*, *CD68*, *SAT2*, *STX8*, *USP43* (x2), *RCVRN* |
| 1q23.3 | 8 | rs2481084, rs117774656, rs1136224, rs11587213, rs16832733, rs11585941, rs11265601, rs12125079 | *F11R*, *UFC1*, *FCER1G* (x2), *PCP4L1*, *C1orf192* (x2), *FCGR2A* |
| 1q22 | 7 | rs79403246, rs3820594, rs17385169, rs3738593, rs1052067, rs2248273, rs16837386 | *GON4L*, *SYT11*, *SSR2*, *MEX3A*, *BGLAP*, *SMG5*, *C1orf61* |
| 7q31.1 | 7 | rs3801936, rs10271385, rs2072208, rs2528650, rs2528651, rs12705437, rs2249956 | *DLD* (x2), *LAMB1* (x5) |
| 19p13.2 | 7 | rs281440, rs17852402, rs75095909, rs13306513, rs5851, rs117129463, rs384148 | *ICAM5* (x3), *LDLR*, *ZNF823* (x2), *ZNF441* |
| 1q21.3 | 6 | rs1879533, rs8480, rs3811414, rs2999551, rs9050, rs12130219 | *CELF3*, *OAZ3*, *TDRKH*, *S100A11*, *TCHH*, *RPTN* |
| 10p14 | 6 | rs7907897, rs7342066, rs7067698, rs10905230, rs12770829, rs7091365 | *KIN* (x4), *ATP5C1*, *GATA3* |
| 19q13.11 | 5 | rs7624, rs7257305, rs17718432, rs3746243, rs2546027 | *ZNF181*, *ZNF599* (x2), *ZNF792* (x2) |
| 7q22.1 | 4 | rs1006507, rs4729676, rs6942609, rs74199090 | *VGF*, *FIS1* (x2), *RABL5* |
| 1p34.1 | 3 | rs56323538, rs882803, rs3748645 | *MUTYH*, *MMACHC*, *NASP* |
| 3p21.1 | 3 | rs4234634, rs74426436, rs2276812 | *NT5DC2*, *SMIM4*, *SPCS1* |
| 4q34.1 | 3 | rs7661281, rs17059534, rs2119788 | *HAND2* (x3) |
| 6p24.3 | 3 | rs3823183, rs16870282, rs9986385 | *DSP*, *TFAP2A* (x2) |
| 11q24.1 | 3 | rs7930974, rs11218912, rs1870761 | *BSX (x3)* |
| 17q21.2 | 3 | rs11079028, rs74647883, rs12386058 | *CNP, NKIRAS2, KCNH4* |
| 1p31.3 | 2 | rs77588186, rs646179 | *L1TD1, USP1* |
| 5p13.3 | 2 | rs17409275, rs6877568 | *DROSHA, C5orf22* |
| 5q14.2 | 2 | rs141671108, rs10045126 | *VCAN (x2)* |
| 6p21.1 | 2 | rs80169011, rs9369458 | *TMEM63B, CAPN11* |
| 7q21.3 | 2 | rs3213654, rs73708843 | *DLX6, DLX5* |
| 7q36.1 | 2 | rs62490272, rs6943752 | *NUB1, RHEB* |
| 9q33.2 | 2 | rs367395, rs2777319 | *STOM, DAB2IP* |
| 10q25.3 | 2 | rs78396600, rs3830026 | *PNLIP, PNLIPRP1* |
| 18p11.31 | 2 | rs1662336, rs7229123 | *MYL12B, TGIF1* |
| 19p13.3 | 2 | rs11085226, rs28384673 | *PTBP1, OAZ1* |
| 19q13.12 | 2 | rs2073900, rs67708190 | *LSR, TMEM147* |
| 19q13.2 | 2 | rs1165838, rs7248468 | *SHKBP1, SNRPA* |
| 1p36.12 | 1 | rs2920 | *ID3* |
| 1p22.1 | 1 | rs1473711 | *GCLM* |
| 2q31.3 | 1 | rs13419200 | *SSFA2* |
| 2q33.1 | 1 | rs6761234 | *SUMO1* |
| 2q36.1 | 1 | rs12694578 | *PAX3* |
| 2q37.2 | 1 | rs6752442 | *GBX2* |
| 3p23 | 1 | rs77280503 | *STT3B* |
| 4p16.2 | 1 | rs10222952 | *STK32B* |
| 5q11.1 | 1 | rs2288468 | *ISL1* |
| 5q14.3 | 1 | rs2290672 | *VCAN* |
| 6q21 | 1 | rs9399964 | *PRDM1* |
| 7q31.31 | 1 | rs2908004 | *WNT16* |
| 9p24.2 | 1 | rs35338539 | *GLIS3* |
| 9p22.3 | 1 | rs13289800 | *NFIB* |
| 10p11.22 | 1 | rs2503997 | *ITGB1* |
| 11p15.5 | 1 | rs6597996 | *DEAF1* |
| 12q13.12 | 1 | rs77349614 | *CSRNP2* |
| 12q22 | 1 | rs11834113 | *SOCS2* |
| 13q14.13 | 1 | rs912784 | *LRRC63* |
| 13q14.2 | 1 | rs79530430 | *MED4* |
| 13q22.2 | 1 | rs41286118 | *UCHL3* |
| 16p11.2 | 1 | rs34286592 | *MAZ* |
| 16q23.1 | 1 | rs749754 | *ADAMTS18* |
| 19q13.31 | 1 | rs892596 | *ZNF180* |
| 20p11.22 | 1 | rs744193 | *NKX2-2* |
| 20q13.12 | 1 | rs80310989 | *KCNS1* |
| 22q11.21 | 1 | rs2531716 | *TANGO2* |

To investigate whether the 137 prioritized SNPs are associated with regulatory activities during craniofacial development, we studied their colocalization with genomic regions harboring H3K4me3 and H3K27ac histone modifications in the epigenomic dataset, which represent active or inducible cis-regulatory elements (26). Around one-half of the SNPs possessed these modifications, of which 37 variants overlapped with H3K4me3 peaks, 17 variants overlapped with H3K27ac peaks, and 26 variants had both histone signatures (S4 Fig). Additionally, 33 of these active SNPs simultaneously possessed repressive histone marks (H3K27me3), indicating that they resided in bivalent “poised” regions, which are known to regulate genes with rapidly alternating expression in stem cells and multipotent progenitors (27). Furthermore, a substantial portion of SNPs not only resided in regions presented with histone peaks, but were also located in proximity to the DEGs of NCCs and cranial placodes, suggesting potential regulatory relationships between the candidate SNPs and these active developmental genes.

### Candidate gene identification and characterization

Our data showed that many of the prioritized SNPs from the machine learning model were associated with active cis-regulatory elements (S3 and S4 Fig). These elements are likely to be involved in gene regulatory networks that control the expression or function of CFM causative genes. Using the SNP-associated cis-regulatory elements, we retrieved their regulatory targets which could be candidate CFM genes by two different approaches (Fig 3A). we first searched for proximal regulatory targets by locating the nearest transcription start site from each SNP using the online tool GREAT (28). Furthermore, we seek distal target genes by examining the topologically associated domain (TAD) where each regulatory element frequently interacts with. The general distribution of these TADs is known to be conserved across different cell types in humans, including progenitors and differentiated cells (29). In addition, functional cis-regulatory elements would contribute to change of gene expression levels. Association between SNP alleles and the expression level (i.e. acting as eQTLs) of another gene suggests interaction between the corresponding element and the gene. Thus, potential distal regulatory targets of the elements were predicted by integrating Hi-C data from H1 human embryonic stem cells and expression quantitative trait loci (eQTL) data from the 3D Genome Browser and Gene-Tissue Expression (GTEx) Portal, respectively (30,31). In particular, genes associated with eQTLs and resided within the same TAD of prioritized SNPs were selected. By combining the two approaches to include both proximal and distal targets, we identified a total of 249 candidate genes for CFM (Fig 3B and S1 Table). The candidate genes were annotated using human and mouse databases (S5 Fig). Notably, comparison with the Human Phenotype Ontology (HPO) and OMIM databases showed that 40 candidates (16%) were known associated genes for CFM, supporting the disease relevance of our findings (Fig 3C and D). Our analysis reveals that many candidate genes for CFM were currently neglected in the HPO and OMIM databases, highlighting the impact of the current study. Among the previously uncharacterized candidates, 11 genes were reported for knockout mouse models with craniofacial phenotypes in the Mouse Genome Informatics (MGI) database (MGI: 0000428) (Fig 3D). A substantial portion of candidates was expressed in NCCs and/or placodal cells according to scRNA-seq data (Fig 3D). Cross-validation with an independent bulk RNA-seq dataset also showed that the candidate genes were expressed at early time points (Carnegie Stage 13 to 17, equivalent to week 4 to 6) (S6 Fig) (13).

**Fig 3.**
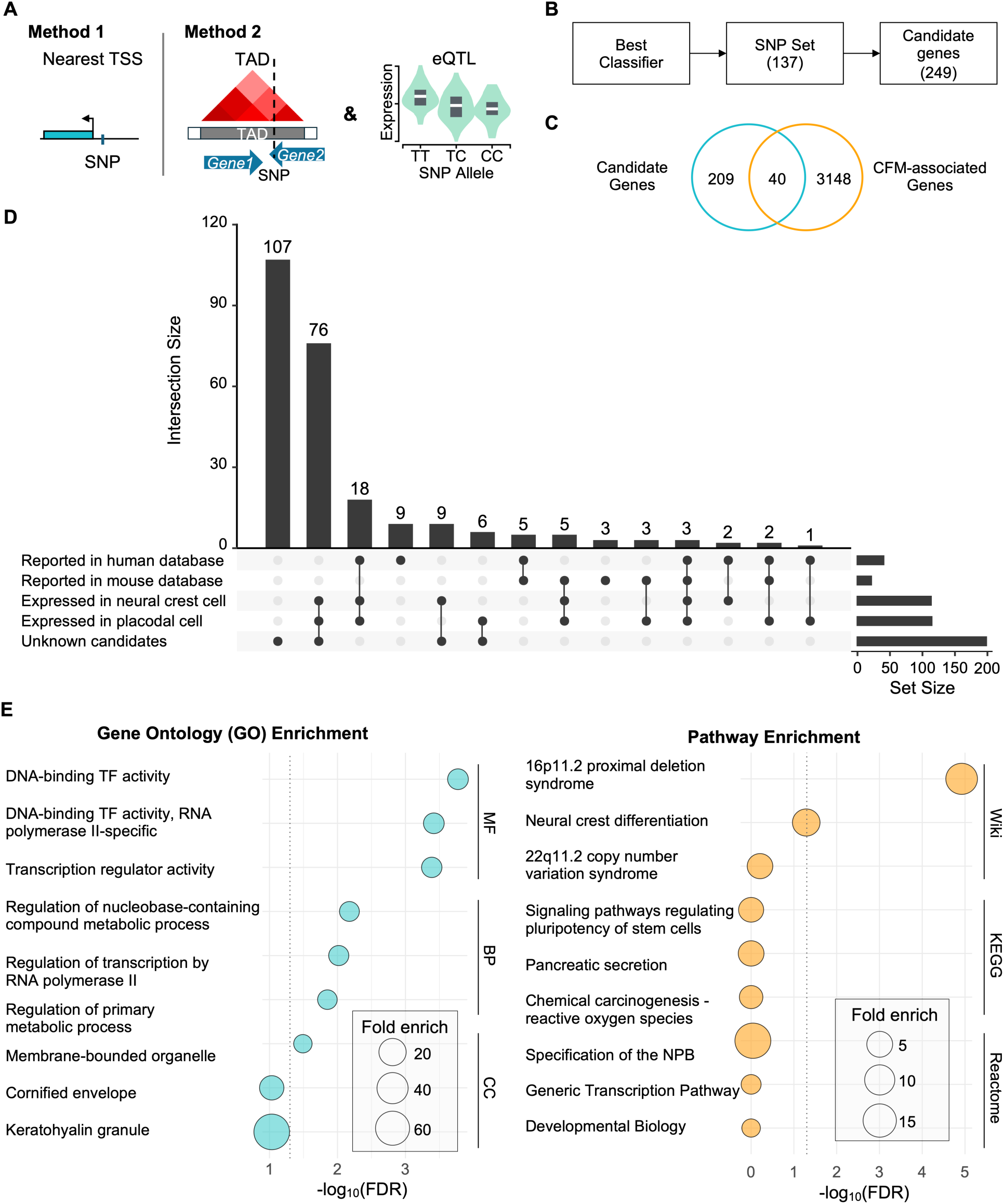
Characterization of the 249 candidate genes associated with prioritized SNPs. (A) Schematic diagram demonstrating the methods used for candidate gene identification from prioritized SNPs. (B) From the SNP set in the best-performing classifier, 249 candidate genes were identified. (C) Venn diagram showing the overlap of candidate genes and reported CFM-associated genes. (D) Upset plot displaying the annotation of candidates. (E) Gene ontology and pathway enrichment results. The top 3 terms of each category are displayed. BP: Biological process; CC: Cellular components; MF: Molecular function; TF: Transcription factor.

Gene ontology (GO) analysis of the candidate genes identified 10, 1, and 14 significantly enriched terms for biological process (BP), cellular components (CC), and molecular function (MF), respectively (Fig 3E, S2 Table). The BP terms fall into 3 categories: regulation of DNA and RNA processes, metabolic processes, and developmental processes. Notably, we observed enrichment of genes related to skeletal system development (GO:0001501), including well-studied transcription factors for development, such as *TFAP2A, DLX5, DLX6, GATA3,* and *HAND2*. Consistently, enriched MF terms were predominantly associated with transcription factor activities. To explore relevant pathways involved by the candidate genes, we performed enrichment analysis using the wikiPathways, KEGG and Reactome databases (Fig3E, S2 Table). Only one wikiPathways term, “16p11.2 proximal deletion syndrome” (Wp4949), was significantly enriched. Patients with 16p11.2 deletion syndrome display intellectual disability, developmental delay and craniofacial dysmorphisms including flat facies, micrognathia, eye and nose abnormalities, and low-set malformed ears (PMID: 29609622, 38605127; OMIM: 611913) (32,33). Interestingly, we observed marginal enrichment of the “Neural crest differentiation” pathway (Wp2064; FDR = 0.0515), revealing that our candidates include not only regulators of late migratory and differentiated neural crest cells (*ITGB1, PAX3, TFAP2A, ISL1, MPZ, MIA*), but also components of the NPB gene regulatory network (GRN) (*GBX2, PAX3, DLX5, TFAP2A*) that act before the specification of neural crest fate. This highlights a potential linkage between early neural crest fate decisions and craniofacial malformations, which has not been extensively explored.

### GRN inference reveals candidates in NCCs and placode regulons

To investigate the role of the candidate genes in neural crest- and placode-specific GRNs, we performed network inference on the ectodermal scRNA-seq data with pySCENIC, which identified 115 regulons (i.e. gene regulatory modules supported by coexpression and motif data) (34,35). Using a regulon specificity score (RSS) cutoff of 0.35, we extracted 10 neural crest- and 16 placode-specific regulons (Fig 4). From the GRNs, we identified 26 distinct transcription factors acting as regulators in neural crest and placode respectively. These regulators were categorized into “NPB Module”, “Neural crest module”, and “Placode modules” based on literature information (36,37). we compared the candidate CFM genes identified in this study with the constructed GRNs. It was observed that 19 of the candidates were target genes in the NCC and placode networks. Among which, 5 genes (*ADAMTS18, DSP, GLIS3, MPZ, TFAP2A*) have been reported as CFM-associated genes in human databases, and 14 genes (*F11R, ISL1, KANK4, L1TD1, LAMB1, MIA, S100A10, S100A11, STOM, STT3B, TESK2, USP43, wDR86, ZNF439*) are novel candidates identified in this study. Interestingly, we discovered 5 candidate CFM genes (*DLX5, DLX6, GATA3, PAX3, PRDM1*) acting as transcriptional regulators in the NPB module, suggesting that some of the CFM genes may have a role in NPB fate decision. Notably, among the candidate genes found inside the NCC and placode GRN, two transcription factors, *PRDM1* and *ISL1,* were supported by other animal functional data (38,39). In the placode-specific network, *PRDM1* is regulated by *GATA3, SIX1, TP63, FOXL2, GRHL2,* and *EMX1*, and was predicted to directly regulate *SIX1, GRHL2, FOXG1, FOXC1, USP43,* and *DSP*. All the regulators and targets of *PRDM1*, except for *EMX1* and *USP43*, are implicated in craniofacial phenotype in human and/or mouse. On the other hand, our results showed that *ISL1* was predicted to be regulated by *GATA3, SIX1, TP63, GRHL2*, and *TBX3* in placodes, all of which are reported to be associated with CFM in human HPO and OMIM databases. Our results strongly suggest that *PRDM1* and *ISL1* are potential causative genes for human CFM.

**Fig 4.**
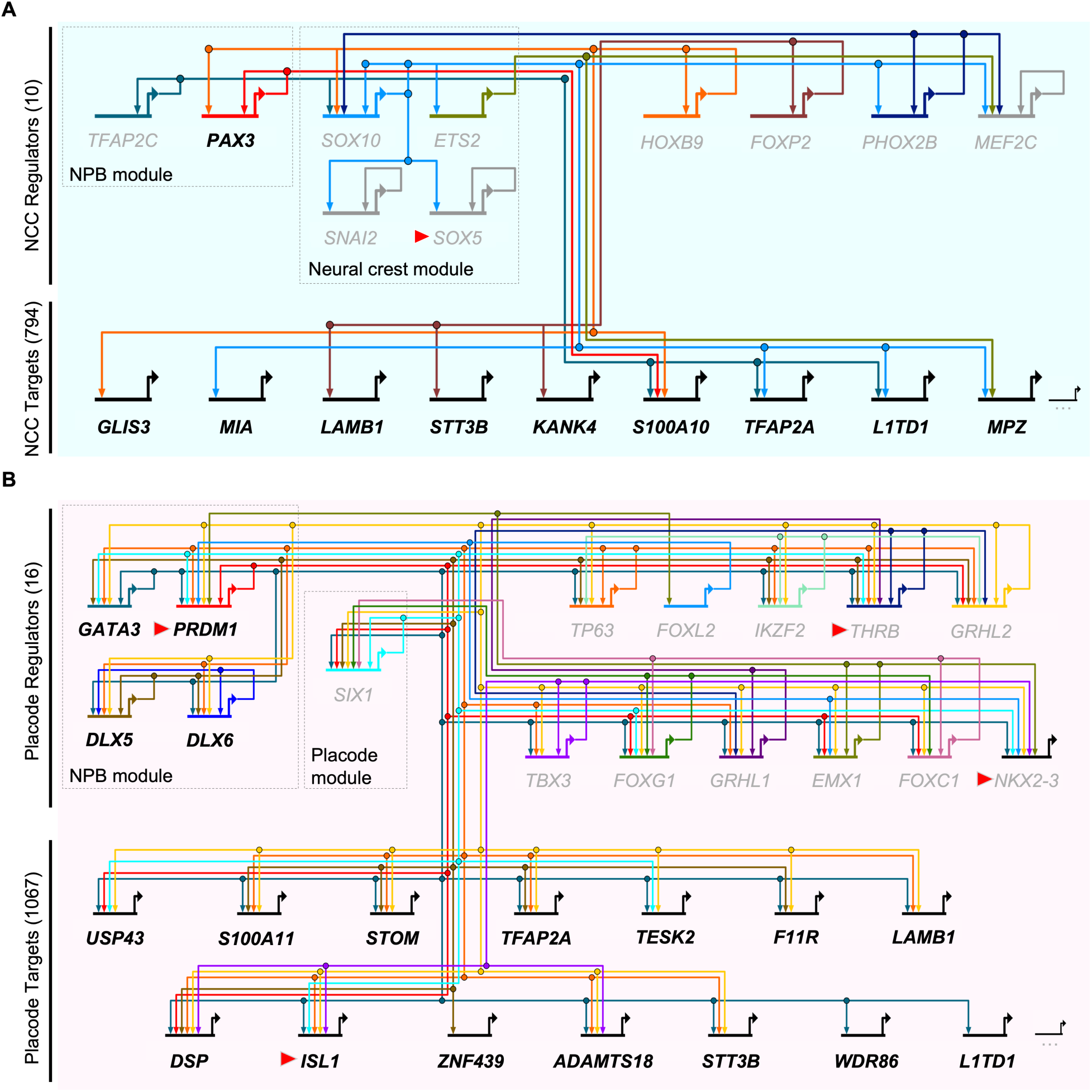
Mapping candidate CFN genes to NCC and placode GRN. (A) BioTapestry plot displaying the network of NCC-specific regulons identified from scRNA-seq of PCW3-4 ectodermal cells. (B) BioTapestry plot showing the network of placode-specific regulons. Transcriptional regulators functioning during neural plate border (NPB), neural crest cell (NCC), and placode differentiation are labelled with grey boxes. Candidate CFM genes identified in this study are indicated in black bold type. CFM genes functionally validated by mutant mouse analyses, but not yet included in human databases are labelled with red arrows.

### Motif analysis identifies candidate functional SNPs for neural crest and placode development

Given that the CFM candidate genes were associated with transcriptional regulation of NCCs and placodal cells (Fig 3B, 4A-B), we investigate whether the alleles of our prioritized SNPs have functional impacts on the neural crest- and placode-expressed transcription factors. Using the R package “motifbreakR”, which assesses the probability of a SNP disrupting transcription factor binding, we discovered four candidate SNPs (rs1662336, rs749754, rs2546027, and rs3820594) not only altered binding motifs and resided in active cis-regulatory elements, but their corresponding upstream transcription factors were also expressed in NCCs and cranial placodes. we also confirmed that the four SNPs were eQTLs for either placode DEGs or known CFM genes using the GTEx database. Two candidates, rs1662336 and rs749754, were located in active chromatin regions near the transcription start sites of MYL12B and ADAMTS18, two placode DEGs, respectively (Fig 5A-B). Particularly, the candidate SNP rs1662336 was predicted to disrupt a *PRDM1* binding motif and acts as an eQTL for *MYL12B*, while the candidate rs749754 was predicted to disrupt the binding site of *SREBF2* and act as an eQTL for *ADAMTS18*. Additionally, we discovered two SNPs, rs2546027 and rs3820594, resided in active cis-regulatory elements and were found to be eQTL for *SCN1B* and *RIT1*, two distal CFM genes (Fig 5C-D). The candidate rs2546027 was predicted to create an ectopic binding site for *GLI3*, while serving as an eQTL for *SCN1B*, a gene associated with cleft palate (HP:0000175). The candidate rs3820594, on the other hand, was predicted to disrupt the binding site of *BACH2*, and act as an eQTL for multiple genes, which including *RIT1*, a gene causing Noonan syndrome (OMIM: 615355). Our results revealed candidate SNPs potentially functional during craniofacial development. This analysis once again identified *PRDM1*, highlighting its likelihood of being a human CFM gene.

**Fig 5.**
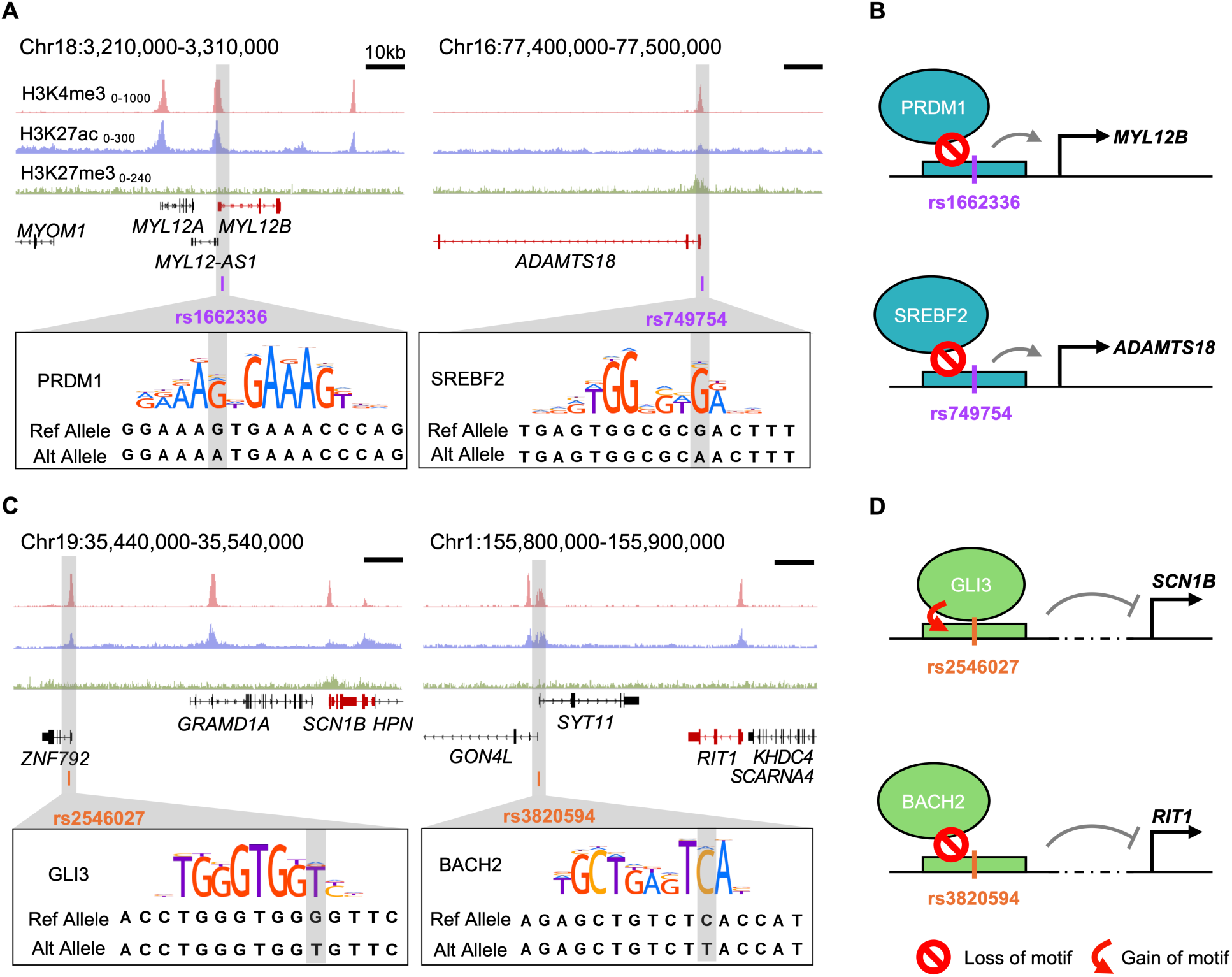
Prioritized SNPs with putative regulatory functions in NCCs and placodes. (A) Predicted regulatory SNPs (purple) disrupting transcription factor binding sites of PRDM1 and SREBF2 respectively. The genome browser tracks show that the SNPs are located in active genomic regions near transcription start sites of their expression quantitative trait loci (eQTL)- associated genes (red). The boxes indicate the disruption of binding motifs by the alternative allele of the prioritized SNPs. (B) Schematic diagrams demonstrating the potential effects of the SNPs on the regulation of their proximal genes. The alternative alleles of rs1662336 and rs749754 disrupt the binding of PRDM1 and SREBF2 to a site near the transcription start sites of *MYL12B* and *ADAMTS18*, affecting their expression levels. (C) Predicted regulatory SNPs (orange) disrupting binding sites of *GLI3* and *BACH2* respectively. The genome tracks shows the location of the eQTL-associated distal genes (red). Disruptions to the binding motifs are indicated by the boxes. (D) Diagrams showing the potential effects of the SNPs on the regulation of their distal genes. The alternative allele of rs2546027 creates an ectopic GLI3 binding site, while the alternative allele of rs3820594 disrupts the binding site of BACH2. These changes affect the expression levels of *SCN1B* and *RIT1*.

## Discussion

In this study, we developed a multiomic approach for disease gene identification by integrating GWAS on craniofacial malformation and functional genomic data on human embryonic tissues, focusing on early developmental time points and two critical progenitor populations, NCCs and cranial placodes. we implemented a weight formula and machine learning classifiers, a type of supervised learning algorithm that predicts binary outcomes from given features. Previous studies have implemented classification models in different manners, such as disease risk assessment and predicting functional SNPs (40,41). Here we applied classifiers to systematically evaluate SNP sets prioritized by different combinations of functional genomic data by their predictive power. Our results showed that the best-performing set of SNPs was prioritized using DEGs of NCCs and cranial placodes, in addition to three histone modification marks. Among the top 20 combinations, the H3K36me3 modification, NCCs, and placodal cells have the highest prevalence. H3K36me3 histone modifications generally represent actively transcribed genes, indicating the contribution of gene expression information to candidate prioritization (26). The involvement of NCCs is consistent with our understanding that they are the major contributors to craniofacial structures and to a wide range of craniofacial disorders. It is reasonable that a substantial portion of craniofacial disease genes can be identified using the NCC expression profile.

Interestingly, the cranial placode transcriptome also facilitated disease gene identification in our model. Placodal defects are implicated in craniofacial abnormalities in human and mouse, yet no previous attempt has incorporated the biology of cranial placodes into CFM gene discovery. Interactions between NCCs and placodal cells are required for proper craniofacial morphogenesis, where the placodes produce signaling molecules to guide NCC behaviors (42,43). In addition to acting as potential signaling epithelia, the placodal cells also directly contribute to the craniofacial structures (15,16,44,45). we previously demonstrated that the *Pax2*-expressed posterior placodal area, apart from the otic and neurogenic epibranchial placodes, also derived into proximal pharyngeal epithelia and epibranchial-derived non-neurogenic domains, which were patterned alternately along the rostro-caudal axis (16). A recent single-cell transcriptomic analysis on human tissues also reported the enrichment of de novo variants of cleft lip with cleft palate in olfactory placode markers, further implicating the linkage between placodal cells and craniofacial abnormalities (19).

We identified 249 candidate genes from the best-performing SNP sets. A portion of the candidates were documented in human databases and were supported by knockout mouse data. Most of the genes were expressed in NCCs and placodal cells and were associated with transcriptional and developmental processes, as indicated by enrichment analyses. Further investigation into the NCC and placode GRN revealed that the predicted regulons overlapped with several CFM candidates, including 15 genes not documented in the HPO and OMIM databases. Among these novel candidates, two transcription factors, *ISL1* and *PRDM1,* were supported by previous animal functional studies (38,39). *ISL1* has been proposed as a candidate for congenital heart disease, but has not been reported as a risk gene for human CFM (46). A recent study showed that epithelial-specific removal of *Isl1* causes cleft palate by disrupting mesenchyme proliferation during palatogenesis (39). *Isl1* is also expressed in the developing olfactory placodes, and the lineages give rise to both sensory and non-sensory epithelial cells (47). *PRDM1* has been associated with Split hand/foot malformation (SHFM), where patients occasionally present craniofacial abnormalities (48). *PRDM1* with SHFM-associated variants failed to rescue craniofacial defects in *prdm1a*-depleted zebrafish in transient overexpression assays, while the wild-type version partially rescued the palatoquadrate, indicating the functional significance of *PRDM1* as a candidate gene for human craniofacial defects (38). Our GRN construction suggests the possibility of *PRDM1* functioning in regulatory network of placode, likely during early NPB specification or differentiation. In agreement with this, *Prdm1* has been demonstrated in chick as a sequential regulator of the neural, neural crest, and placodal lineages, pointing to a linkage between progenitor specification and craniofacial abnormalities (49). Our regulatory SNP analysis also independently discovered *PRDM1*. In particular, we predicted that the variant rs1662336 may disrupt a *PRDM1* motif and acts as an eQTL for another placode DEG, *MYL12B*. *MYL12B* is known for regulating actin filament organization, yet its function in placodes or other epithelia is unexplored (50).

Taken together, we demonstrated a method that successfully integrated multiomic information for candidate gene prioritization, focusing on specific cell types during early development. Our approach can be easily extended to different cell types, time points, and other congenital disorders, or be scaled up with a larger sample size and a more advanced machine learning model. we also provided a list of candidate genes and several putative gene regulations underlying human craniofacial malformation. More importantly, our results suggested potential links between craniofacial malformation with placodal cells and NPB specification, which has not been extensively investigated in previous studies. This study provides new perspectives on the cellular mechanisms and pathogenic process of CFM.

## Materials and Methods

### SNP array collection and processing

A collection of SNP array datasets was downloaded from the Gene Expression Omnibus (GEO) (Accession: GSE69664, GSE141901, GSE74100, and GSE248483) (6,51,52). These assays were conducted in the Asian population and involved 939 patients from China in addition to 447 healthy individuals from China, Myanmar, and Vietnam. Raw SNP array data were downloaded in the IDAT format and processed with the genotyping module of Illumina GenomeStudio 2.0 using a genotype call threshold of 0.15. To ensure reliable genotype calls, we filter out SNPs with GenTrain score below 0.7, which measures the quality of the SNP clustering process in GenomeStudio. Subsequent quality control process was performed with PLINK1.9, where we selected variants with SNP call rate >95%, sample call rate >80%, and minor allele frequencies >1%, and had a Hardy-weinberg equilibrium score >1e-4. Phasing and genotype imputation were conducted with Beagle5.4 using 1000 Genomes Project Phase 3 (Genome build 37) as a reference panel after merging the datasets (53). To filter out low-quality imputation results, we set a threshold of DR2 > 0.8. LD pruning was then carried out (--indep-pairwise 200kb 1 0.5), which removes variants with R2 >0.5, and regions with high LD were trimmed (54,55). As a result, 161,092 variants passed quality control. The dataset was subsequently separated into two subsets for the association test and building logistic regression classifiers. Association test for disease was performed on 493 patients and 207 controls with PLINK1.9 using the logistic model and the first 10 principal components (PCs) as covariates to control for population stratification.

### scRNA-seq data processing

scRNA-seq datasets of post-conceptual week (PCW) 3 and 4 human embryonic heads were retrieved from GEO (Accession: GSE155121) (20). Raw FASTQ files were downloaded and aligned to the reference genome (hg19) using Cell Ranger (v7.1.0) (56). Quality control was performed with scanpy (v1.9), where cells with >200 expressed genes, unique gene counts >1,000 and <10,000, total counts >1,000 and <70,000, and mitochondrial counts <60% were kept, resulting in an AnnData object with 49,123 cells and 36,601 features (57). Dimension reduction was conducted after data normalization, log-transformation, and feature selection, and the first 10 principal components were used for constructing the nearest neighbor graph, which was visualized with the Uniform and Manifold Approximation and Projection plot. Cell clustering was done with the Leiden method, and ectodermal derivatives were isolated for subsequent analysis. The Python package Palantir was used to compute a diffusion space for the ectodermal cells and generated a 2-dimensional embedding using the Force Atlas 2 algorithm (22). Subpopulations were annotated with cellular markers. Differentially expressed genes (DEGs) were identified using the FindAllMarkers function in Seurat (v4) with a minimum log-fold change >0.25 and expression detected in >25% of cells. Ribosomal genes are excluded from the DEG list. The top 10,000 DEGs of targeted cell types were used for downstream variants prioritization.

### ChIP-Seq data processing

Histone modification ChIP-Seq data from developing human craniofacial tissues were downloaded from GEO via the accession code GSE97752 (12). we selected data between PCW 4 and 5 (Carnegie stage 13, 14, 15) to match the developmental time points of the scRNA-Seq data. TagAlign files were extracted and converted to BAM format using the bedtobam function from BEDTools (v2.31.0) (58). The resulting BAM files were sorted with samtools (v1.6), and index files were created (59). MACS2 (v2.2.9.1) was then employed for peak calling using a Q-value threshold of 0.01 (60). Downstream analyses were performed using the average peak scores for each histone modification type.

### Bulk RNA-Seq data processing

Expression matrix of bulk RNA-Seq data was downloaded from http://cotneyweb.cam.uchc.edu/craniofacial_bulkrna/ (13). Z-score calculation and hierarchical clustering were performed with R (v4.4.0).

### Functional score

To prioritize variants, we designed a weighted formula to calculate the functional score (FS), which is computed by multiplying the negative logarithm of the association test p-value by the product of weights derived from additional functional genomic data:

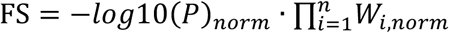

For scRNA-seq data, the DEGs in cell populations of interest were identified, and the corresponding wilcoxon Rank Sum test p-values were used to approximate marker gene specificity and applied as weights. For epigenomic data, average peak scores were used as weights. SNPs that fall within ±400kb from the DEG, or inside the ChIP-Seq peaks, were identified using the intersect function from bedtools (v2.31.0), and assigned with the corresponding weight values. The negative logarithm of p-value and weights were scaled to 0-100, followed by the addition of a pseudocount of 1.

### Combinatorial analysis

Logistic regression classifiers were used to identify the combinations of additional functional genomic data that improve disease classification power when they were implemented in the functional score calculation. In particular, all possible combinations are generated from the fixed set of transcriptomic and epigenomic data (number of combinations 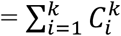). FS was calculated iteratively for each combination, where top SNPs were retrieved for subsequent analysis. To construct a logistic regression classification model, SNP array data of 446 craniofacial malformation patients and 240 controls were used as a sample for model training and testing. These data were assayed by the same array platform (Illumina HumanOmniZhongHua-8 v1.0 Beadchip) and independent of the earlier dataset used for association test, to reduce the risk of batch effect and data leakage. To avoid overfitting, we included the top 137 variants in each SNP set as features, maintaining a 1:5 feature-to-sample ratio. To assess the performance of the logistic regression classifiers, the samples were randomly split into subsets with a 7:3 ratio, where 70% of the data was used for training and the remaining 30% for testing. The process was repeated for 200 iterations to reduce variability due to random splits. In each iteration, the trained model was evaluated on the test set using the receiver operating characteristic (ROC) curve as an evaluation metric, where the area under the ROC curve (AUROC) was employed to quantify the classification power of the model. The average AUROC of 200 iterations was used to compare the power of different classifiers.

### Enrichment analysis

Functional enrichment analysis of candidate genes was performed using the Database for Annotation, Visualization, and Integrated Discovery (DAVID) online tool (61). Enrichment was assessed for the GO categories, including Biological Process, Molecular Function, and Cellular Components (62). Pathway enrichment analyses were performed with the wikiPathways, KEGG, and Reactome databases (63–65).

### GRN inference

GRN reconstruction was performed on scRNA-seq data using pySCENIC (v0.12.1) (34). Regulons were computed using default parameters. RSS were computed across all annotated cell types. Cell type-specific regulons were identified using a cutoff of RSS = 0.35.

### Motif analysis

To predict the effect of SNPs on gain or loss of transcription factor binding motifs, we implemented the R package motifbreakR using default parameters and SNP information from the dbSNP Build 155 (66). For motif data, we applied the core collections of HOCOMOCO (v11) (67). Candidate SNPs predicted to have strong disruptions on motifs were identified.

### Database search

The NHGRI-EBI GWAS Catalog was used to retrieve genomic loci associated with craniofacial microsomia (Experimental Factor Ontology (EFO) ID: MONDO_0015397), outer ear morphological trait (EFO ID: EFO_0007664) and cleft lip with/without cleft palate (EFO ID: EFO_0003959). (25). Information from the HPO and OMIM databases were implemented to identify documented genes associated with craniofacial malformation (68,69). For HPO, we extracted all associated genes under the category “Abnormality of the face” (HP:0000271). For OMIM, we obtained the gene list curated by Naqvi et. al, where they manually screened all OMIM records using the query “craniofacial” and extracted only genes actually related to craniofacial malformation (2). To collect knockout mouse information related to craniofacial abnormalities, we identified genes under the “abnormal craniofacial morphology” (MP:0000428) category in the MGI database (70). eQTL data and TADs in H1 human embryonic stem cells were retrieved from the GTEx portal and 3D Genome Browser, respectively, for candidate gene identification (30,31). The RegulomeDB and FORGEdb were implemented during SNP annotation to assess functional potentials (23,24).

### Statistical test

All numerical data were analyzed statistically with student’s t-test for comparing different groups using R (v4.4.0). A probability of P < 0.05 will be considered significant.

## Acknowledgements

We thank Dr. Matthew Sai Pong Ho and Dr. Mengmeng Shi for helpful discussions. This project is supported by a research grant from the Health and Medical Research Fund (HMRF 08191246) to MHS.

---

**S1 Fig.**
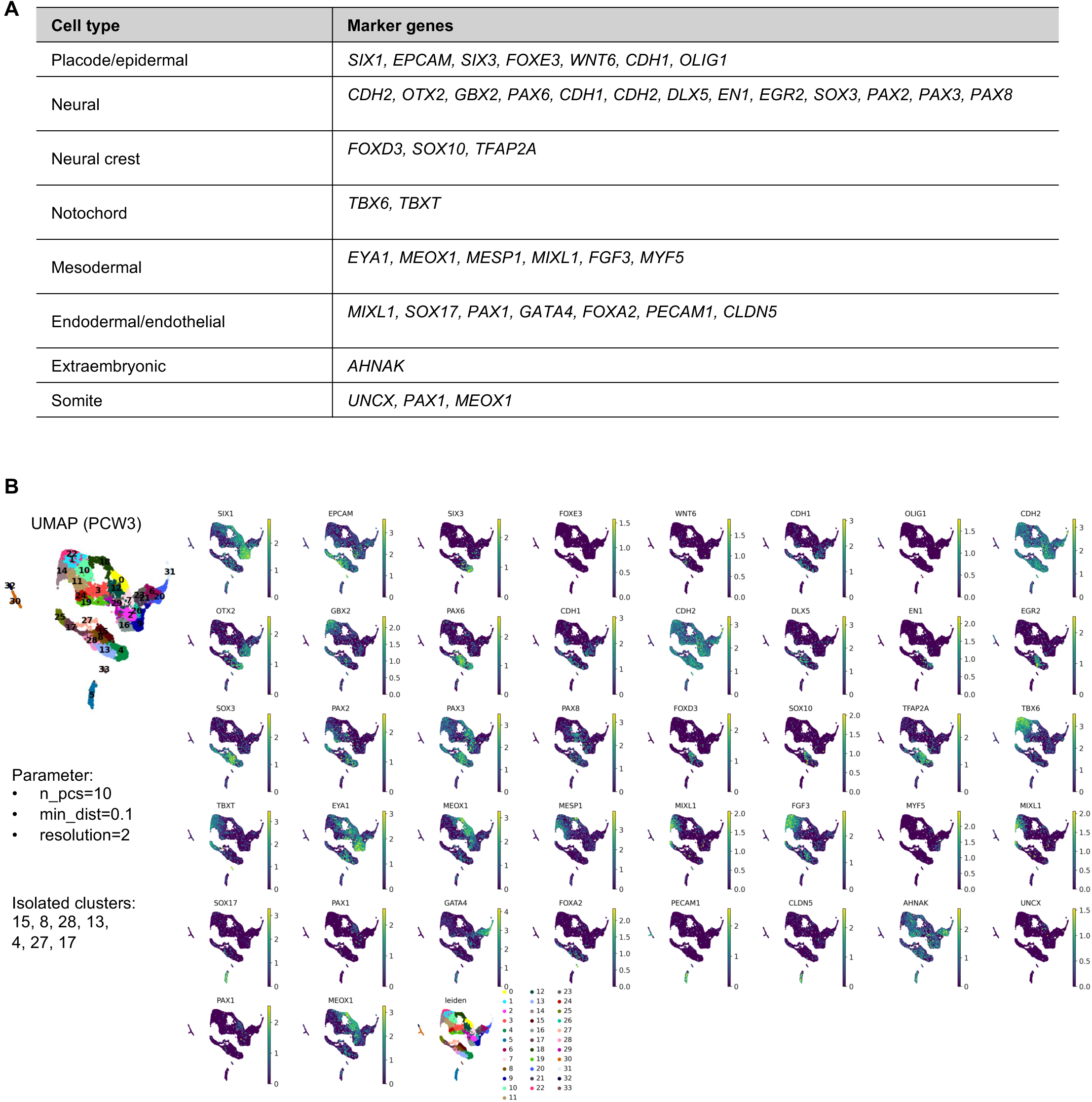
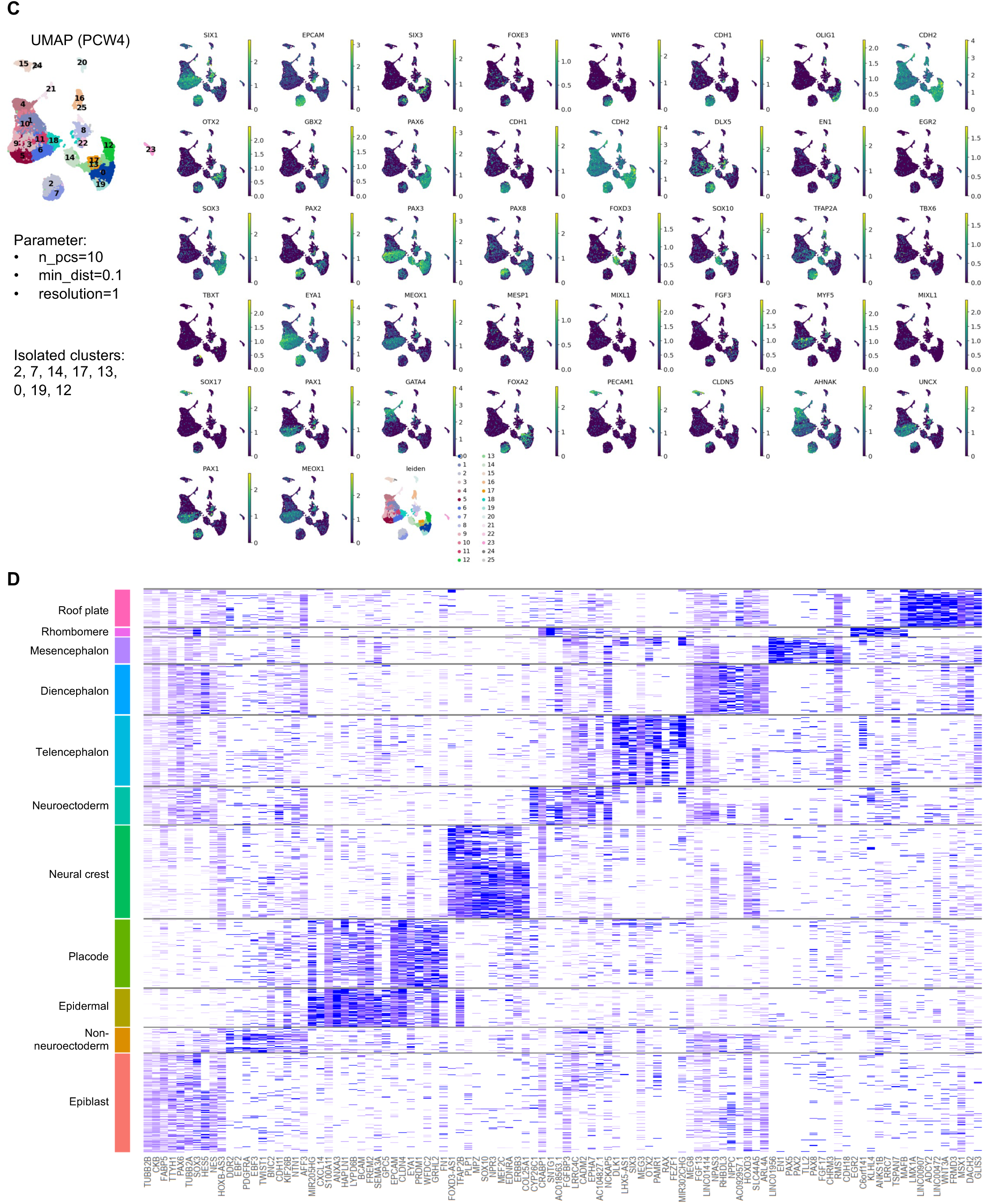
Annotation of scRNA-seq data. (A) Marker genes for annotation of PCW3 and PCW4 human embryo scRNA-seq. (B-C) Parameters for dimension reduction and clustering, as well as ectodermal Leiden clusters isolated for downstream analysis were shown on left. Feature plots on right display expression pattern of marker genes. (D) Heatmap showing top marker genes of 11 annotated ectodermal clusters isolated from (B) and (C).

**S2 Fig.**
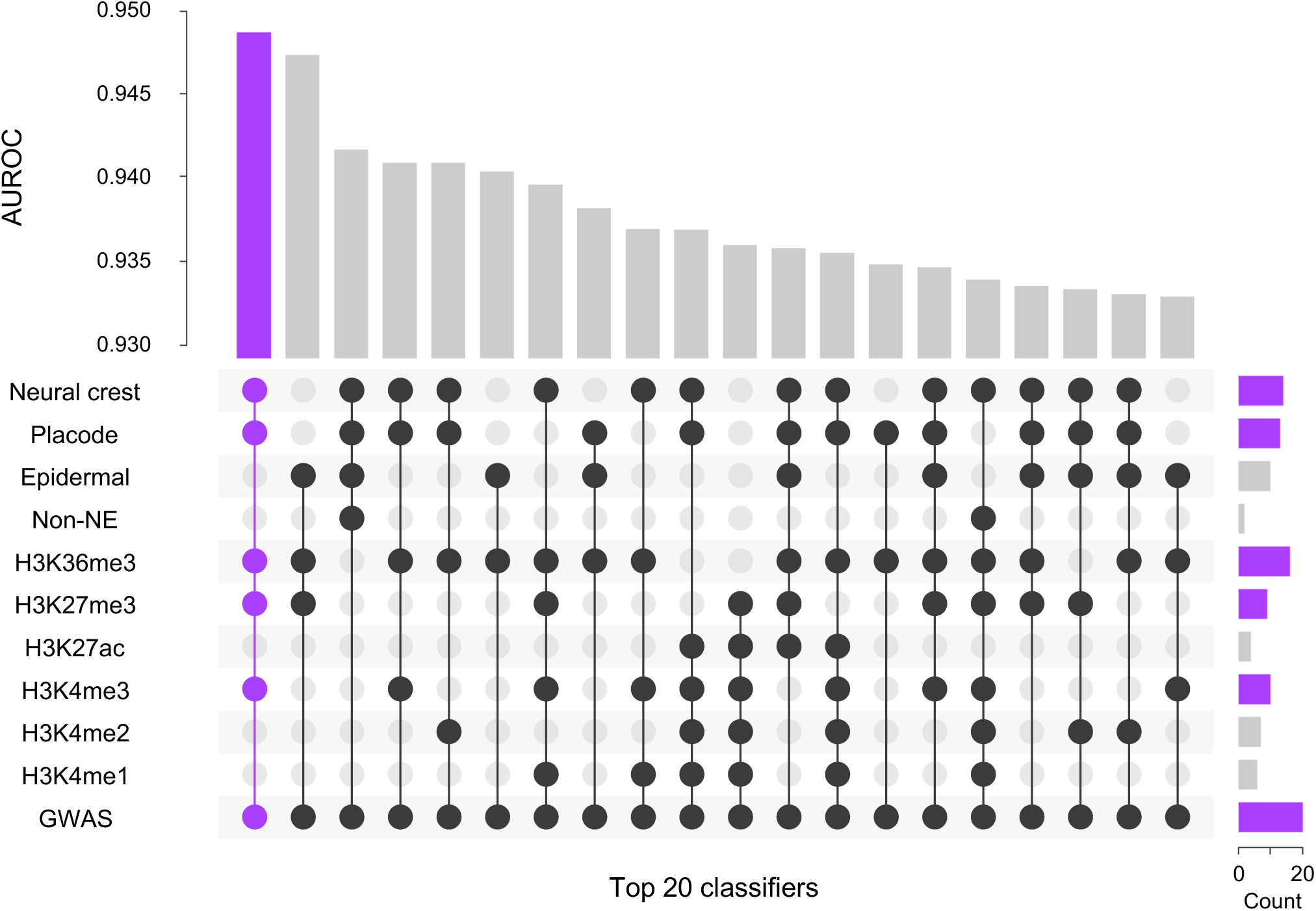
Combinations of data layers in the top 20 machine learning classifiers. Upset plot displayed the combination of data layers included in the top 20 classifier. Bar at top visualized the average AUROC. Bar on right shows the frequency of the data layer shown on plot.

**S3 Fig.**
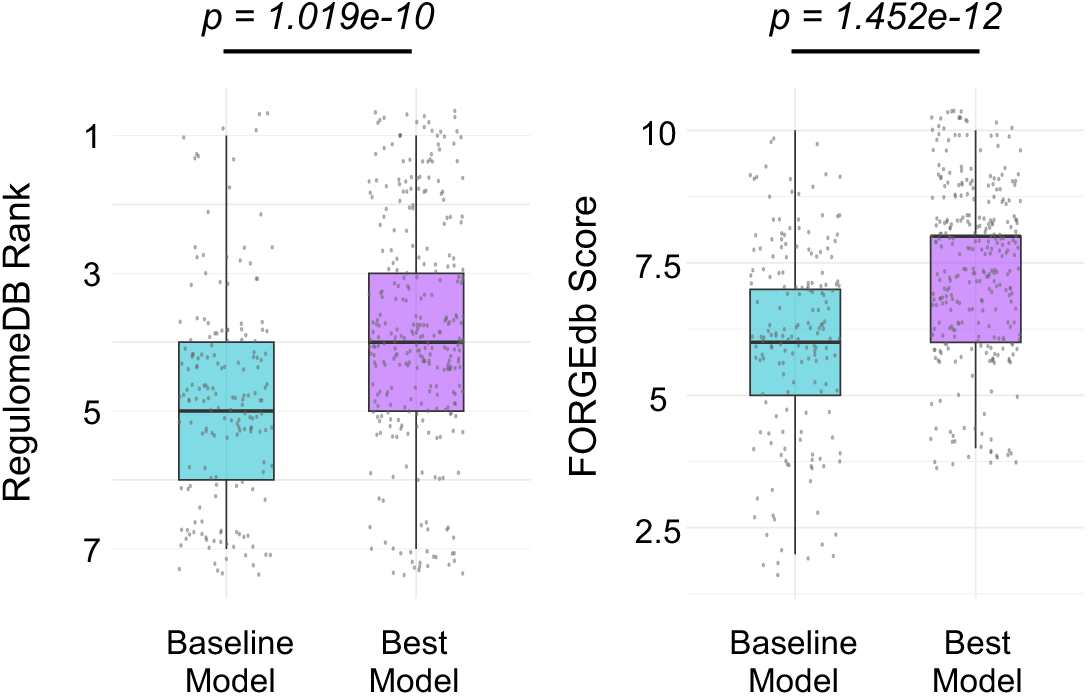
Regulatory scores of SNP sets in best-performing and baseline classifiers. Box plots comparing the RegulomeDB rank and FORGEdb scores of the SNP sets from the best-performing and baseline classifiers.

**S4 Fig.**
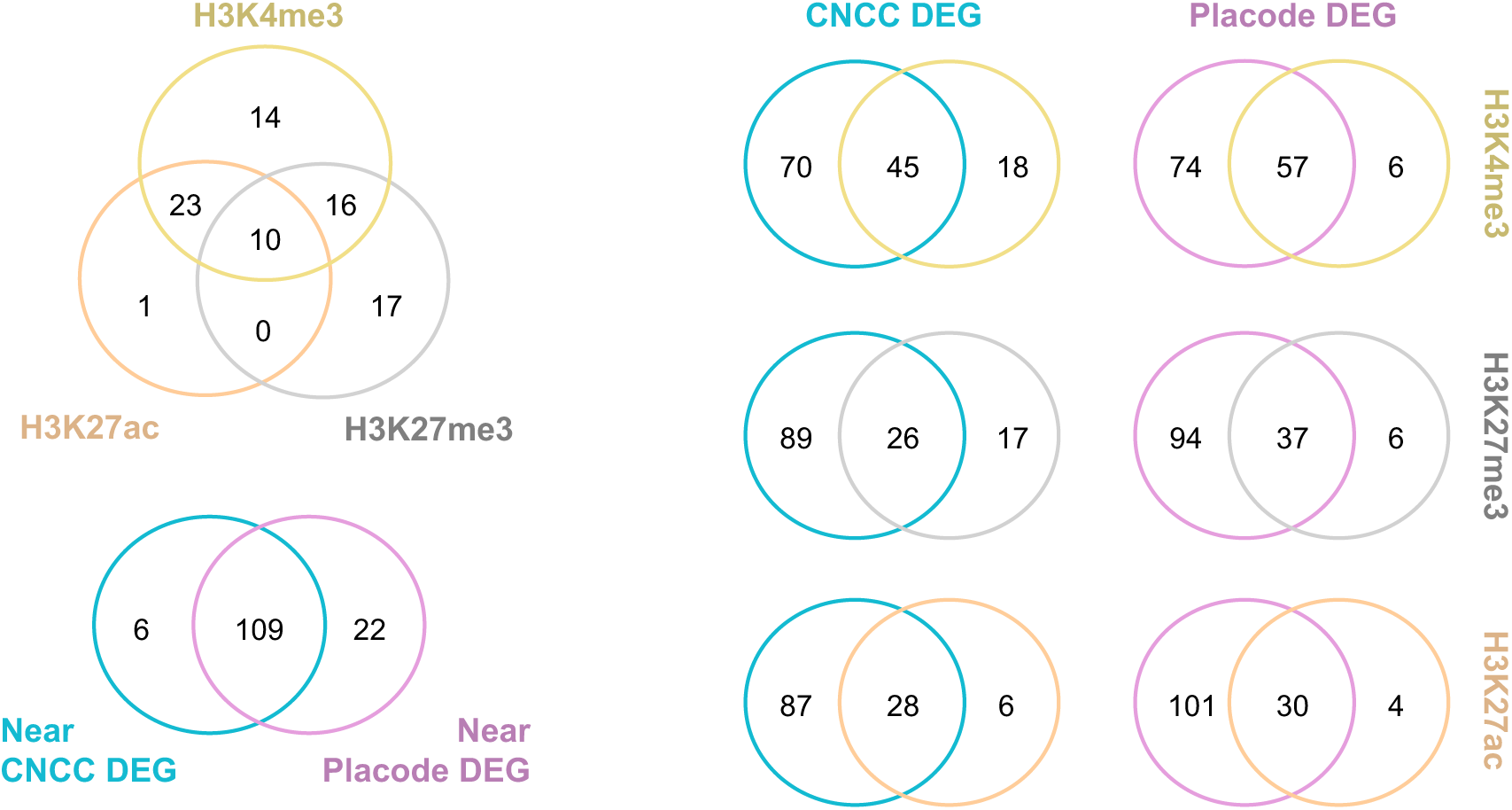
Epigenomic features of prioritized variants. Venn diagram illustrating the number of SNPs located inside H3K4me3/H3K27ac/H3K27me3 peaks and SNPs located ±400kb from DEGs of NCCs or placodes. NCC: Neural crest cell; DEG: differentially expressed gene.

**S5 Fig.**
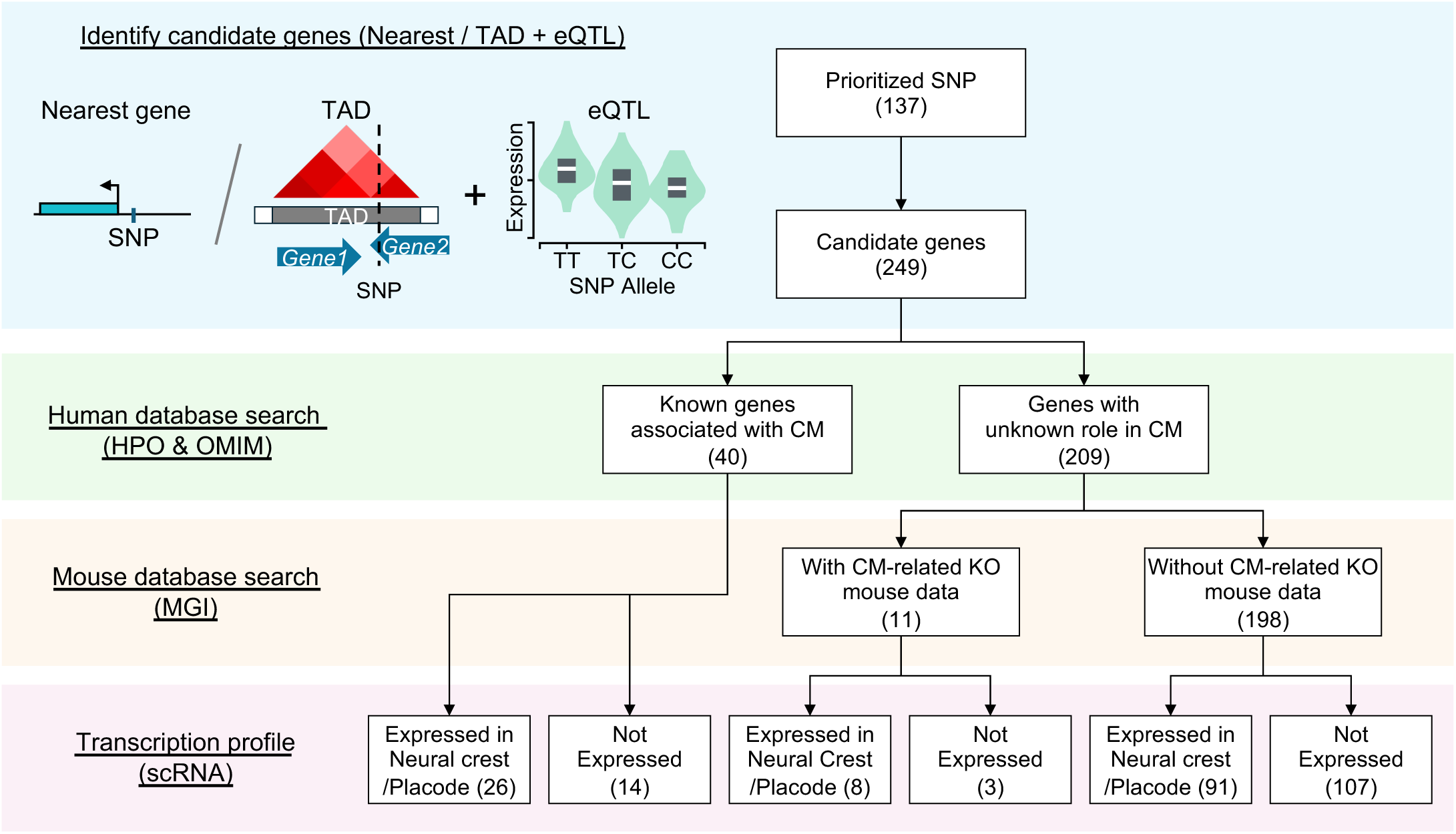
Annotation of 259 candidate genes. Flowchart illustrating the annotation of candidate genes using human/mouse databases and single-cell transcriptome. A gene is considered expressed when its transcript count is higher than 75% of all genes in neural crest cells (NCCs) or placodal cells.

**S6 Fig.**
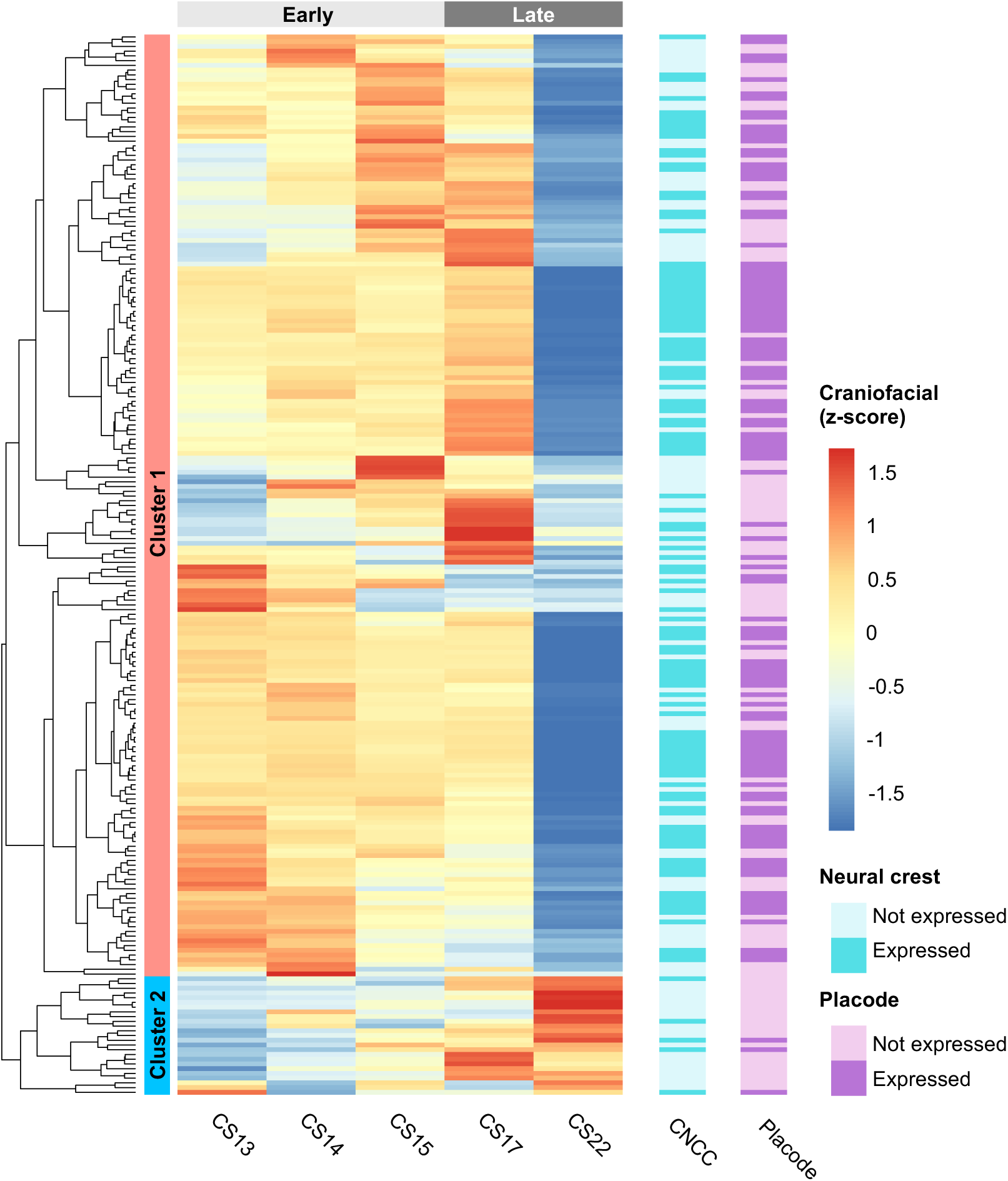
Craniofacial expression profile of 249 candidate genes. Analysis on an independent bulk RNA-seq dataset of developing human craniofacial tissues, comprising three early (PCW 4-5) and two later (PCW 6-8) time points. Heatmap showing expression profile of candidate genes in the dataset. Dendrogram (left) shows result of hierarchical clustering. Cyan and purple bar (right) highlights genes expressed by NCCs and placodes in original scRNA-seq data. A gene is considered expressed when its transcript count is higher than 75% of all genes in neural crest cells (NCCs) or placodal cells.

**S1 Table.** List of 249 candidate genes identified from prioritized SNPs. Candidate genes were identified by two distinct approaches from prioritized SNPs. Potential proximal targets of SNPs were retrieved by locating the nearest transcription start site from each SNP using the online tool GREAT. Potential distal targets were identified by combining eQTL and TAD information.

| SNP | Chr | Position | Functional Score | Locus | Locus Coordinate | Candidate genes |
| --- | --- | --- | --- | --- | --- | --- |
| rs2920 | 1 | 23884780 | 90604.41333 | 1p36.12 | chr1:20400000-23900000 | ASAP3, ID3, ZNF436-AS1, TCEA3 |
| rs56323538 | 1 | 45810797 | 140597.17 | 1p34.1 | chr1:44100000-46800000 | MUTYH |
| rs882803 | 1 | 45976147 | 137467.0572 | 1p34.1 | chr1:44100000-46800000 | CCDC163, MUTYH, PRDX1, TESK2, MMACHC, AKR1A1 |
| rs3748645 | 1 | 46035470 | 111421.4432 | 1p34.1 | chr1:44100000-46800000 | NASP |
| rs77588186 | 1 | 62627726 | 79622.59373 | 1p31.3 | chr1:61300000-68900000 | L1TD1 |
| rs646179 | 1 | 62902874 | 1158411.008 | 1p31.3 | chr1:61300000-68900000 | KANK4, USP1, DOCK7 |
| rs1473711 | 1 | 94385591 | 79341.00653 | 1p22.1 | chr1:92000000-94700000 | GCLM |
| rs1879533 | 1 | 151694241 | 380801.4082 | 1q21.3 | chr1:150300000-155000000 | LINGO4, TUFT1, CELF3, RIIAD1, TDRKH-AS1, MRPL9, SNX27, OAZ3, TDRKH, HRNR |
| rs8480 | 1 | 151733335 | 89687.58594 | 1q21.3 | chr1:150300000-155000000 | RORC, LINGO4, MRPL9, SNX27, OAZ3, TDRKH |
| rs3811414 | 1 | 151763246 | 95603.39795 | 1q21.3 | chr1:150300000-155000000 | LINGO4, TUFT1, CELF3, RIIAD1, TDRKH-AS1, S100A10, MRPL9, SNX27, OAZ3, TDRKH, HRNR |
| rs2999551 | 1 | 152008309 | 146063.6432 | 1q21.3 | chr1:150300000-155000000 | S100A10, THEM4, FLG, S100A11 |
| rs9050 | 1 | 152079314 | 628766.3864 | 1q21.3 | chr1:150300000-155000000 | THEM4, TCHH, LINGO4, C2CD4D-AS1, FLG2, FLG, RIIAD1 |
| rs12130219 | 1 | 152162106 | 192512.9433 | 1q21.3 | chr1:150300000-155000000 | FLG, RPTN, FLG2, CRNN, HRNR |
| rs79403246 | 1 | 155822914 | 80349.88136 | 1q22 | chr1:155000000-156500000 | GON4L |
| rs3820594 | 1 | 155829511 | 89827.08442 | 1q22 | chr1:155000000-156500000 | MSTO2P, SYT11, FDPS, DAP3, MSTO1, PKLR, ASH1L, HCN3, MUC1, ARHGEF2, RIT1 |
| rs17385169 | 1 | 155983869 | 137173.9509 | 1q22 | chr1:155000000-156500000 | RIT1, SSR2, ARHGEF2-AS2, UBQLN4 |
| rs3738593 | 1 | 156045973 | 85571.23497 | 1q22 | chr1:155000000-156500000 | LAMTOR2, SLC25A44, SEMA4A, MEX3A |
| rs1052067 | 1 | 156206121 | 138391.2888 | 1q22 | chr1:155000000-156500000 | SLC25A44, RUSC1-AS1, PMF1, SEMA4A, BGLAP, PAQR6, GLMP, CCT3, SMG5, TMEM79, TSACC, MEF2D |
| rs2248273 | 1 | 156247907 | 133222.7789 | 1q22 | chr1:155000000-156500000 | SEMA4A, PMF1, SLC25A44, BGLAP, PAQR6, GLMP, CCT3, SMG5, TMEM79 |
| rs16837386 | 1 | 156395744 | 107642.2687 | 1q22 | chr1:155000000-156500000 | C1orf61, RHBG |
| rs2481084 | 1 | 160989614 | 129128.2367 | 1q23.3 | chr1:160500000-165500000 | ARHGAP30, ATP1A4, USF1, TSTD1, F11R |
| rs117774656 | 1 | 161123212 | 235181.276 | 1q23.3 | chr1:160500000-165500000 | UFC1 |
| rs1136224 | 1 | 161184097 | 370529.8051 | 1q23.3 | chr1:160500000-165500000 | B4GALT3, FCER1G, ADAMTS4 |
| rs11587213 | 1 | 161184875 | 80697.21191 | 1q23.3 | chr1:160500000-165500000 | FCER1G, PFDN2, KLHDC9, NIT1, TSTD1, F11R, ADAMTS4, B4GALT3, ITLN2, DEDD, NDUFS2, PPOX |
| rs16832733 | 1 | 161228843 | 2705730.427 | 1q23.3 | chr1:160500000-165500000 | TOMM40L, ADAMTS4, B4GALT3, PCP4L1, NDUFS2 |
| rs11585941 | 1 | 161312656 | 82829.17648 | 1q23.3 | chr1:160500000-165500000 | CFAP126, C1orf192, SDHC, MPZ |
| rs11265601 | 1 | 161398984 | 361545.7289 | 1q23.3 | chr1:160500000-165500000 | SDHC, C1orf192, TSTD1 |
| rs12125079 | 1 | 161408556 | 242292.8619 | 1q23.3 | chr1:160500000-165500000 | FCGR2A |
| rs13419200 | 2 | 182758257 | 394411.3251 | 2q31.3 | chr2:180600000-183000000 | SSFA2 |
| rs6761234 | 2 | 203102345 | 120730.2573 | 2q33.1 | chr2:197400000-203300000 | KIAA2012, SUMO1 |
| rs12694578 | 2 | 223161624 | 713338.0942 | 2q36.1 | chr2:221500000-225200000 | PAX3, CT75 |
| rs6752442 | 2 | 237073776 | 109004.9214 | 2q37.2 | chr2:235600000-237300000 | ASB18, GBX2 |
| rs77280503 | 3 | 31575382 | 85246.31386 | 3p23 | chr3:30900000-32100000 | STT3B |
| rs4234634 | 3 | 52557069 | 561612.7369 | 3p21.1 | chr3:52300000-54400000 | NT5DC2 |
| rs74426436 | 3 | 52611711 | 133957.8922 | 3p21.1 | chr3:52300000-54400000 | SMIM4 |
| rs2276812 | 3 | 52729960 | 239127.6327 | 3p21.1 | chr3:52300000-54400000 | SPCS1 |
| rs10222952 | 4 | 5053674 | 173209.1965 | 4p16.2 | chr4:4500000-6000000 | CYTL1, STK32B |
| rs7661281 | 4 | 174437070 | 85117.41213 | 4q34.1 | chr4:171900000-176300000 | HAND2, HAND2-AS1 |
| rs17059534 | 4 | 174444902 | 1266149.65 | 4q34.1 | chr4:171900000-176300000 | HAND2 |
| rs2119788 | 4 | 174447389 | 1049882.991 | 4q34.1 | chr4:171900000-176300000 | HAND2, HAND2-AS1 |
| rs17409275 | 5 | 31514127 | 350663.3706 | 5p13.3 | chr5:28900000-33800000 | DROSHA |
| rs6877568 | 5 | 31532685 | 125461.3898 | 5p13.3 | chr5:28900000-33800000 | C5orf22 |
| rs2288468 | 5 | 50683655 | 94655.95722 | 5q11.1 | chr5:48400000-50700000 | ISL1 |
| rs141671108 | 5 | 82772272 | 89044.26711 | 5q14.2 | chr5:81400000-82800000 | VCAN |
| rs10045126 | 5 | 82782034 | 102291.7556 | 5q14.2 | chr5:81400000-82800000 | VCAN |
| rs2290672 | 5 | 82850587 | 103843.2016 | 5q14.3 | chr5:82800000-92300000 | VCAN-AS1, VCAN |
| rs3823183 | 6 | 7543468 | 343204.9046 | 6p24.3 | chr6:7100000-10600000 | DSP |
| rs16870282 | 6 | 10422524 | 181893.9931 | 6p24.3 | chr6:7100000-10600000 | TFAP2A |
| rs9986385 | 6 | 10424933 | 228037.0166 | 6p24.3 | chr6:7100000-10600000 | TFAP2A |
| rs80169011 | 6 | 43969839 | 817592.1913 | 6p21.1 | chr6:40500000-46200000 | TMEM63B |
| rs9369458 | 6 | 44116993 | 78524.88402 | 6p21.1 | chr6:40500000-46200000 | MRPL14, CAPN11, TMEM63B |
| rs9399964 | 6 | 106441922 | 563146.6153 | 6q21 | chr6:105500000-114600000 | PRDM1 |
| rs3213654 | 7 | 96639368 | 1557956.095 | 7q21.3 | chr7:92800000-98000000 | DLX6 |
| rs73708843 | 7 | 96654349 | 161060.8189 | 7q21.3 | chr7:92800000-98000000 | DLX5 |
| rs1006507 | 7 | 100814034 | 169540.8482 | 7q22.1 | chr7:98000000-103800000 | VGF |
| rs4729676 | 7 | 100894697 | 76330.70931 | 7q22.1 | chr7:98000000-103800000 | FIS1 |
| rs6942609 | 7 | 100895480 | 134514.5646 | 7q22.1 | chr7:98000000-103800000 | FIS1 |
| rs74199090 | 7 | 100959859 | 83907.37363 | 7q22.1 | chr7:98000000-103800000 | RABL5 |
| rs3801936 | 7 | 107536417 | 88284.69371 | 7q31.1 | chr7:107400000-114600000 | DLD, CBLL1, CBLL1-AS1, SLC26A3 |
| rs10271385 | 7 | 107556623 | 145299.9768 | 7q31.1 | chr7:107400000-114600000 | DLD, CBLL1, CBLL1-AS1, SLC26A3 |
| rs2072208 | 7 | 107591914 | 259237.9407 | 7q31.1 | chr7:107400000-114600000 | DLD, LAMB1, CBLL1 |
| rs2528650 | 7 | 107600558 | 140798.0326 | 7q31.1 | chr7:107400000-114600000 | LAMB1, DLD |
| rs2528651 | 7 | 107602624 | 125152.9441 | 7q31.1 | chr7:107400000-114600000 | LAMB1 |
| rs12705437 | 7 | 107633517 | 175047.3928 | 7q31.1 | chr7:107400000-114600000 | LAMB1, DLD |
| rs2249956 | 7 | 107641616 | 78813.90469 | 7q31.1 | chr7:107400000-114600000 | LAMB1 |
| rs2908004 | 7 | 120969769 | 87129.38935 | 7q31.31 | chr7:117400000-121100000 | WNT16, FAM3C |
| rs62490272 | 7 | 151039179 | 159692.5561 | 7q36.1 | chr7:147900000-152600000 | NUB1 |
| rs6943752 | 7 | 151215950 | 110731.001 | 7q36.1 | chr7:147900000-152600000 | WDR86, RHEB |
| rs35338539 | 9 | 4298589 | 186633.1533 | 9p24.2 | chr9:2200000-4600000 | GLIS3 |
| rs13289800 | 9 | 14348827 | 76408.55105 | 9p22.3 | chr9:14200000-16600000 | NFIB |
| rs367395 | 9 | 124131512 | 135586.1481 | 9q33.2 | chr9:122500000-125800000 | RAB14, GSN-AS1, STOM |
| rs2777319 | 9 | 124362502 | 92122.74129 | 9q33.2 | chr9:122500000-125800000 | DAB2IP |
| rs7907897 | 10 | 7794648 | 328845.5306 | 10p14 | chr10:6600000-12200000 | KIN, ITIH2, TAF3, ATP5F1C |
| rs7342066 | 10 | 7795911 | 188617.8166 | 10p14 | chr10:6600000-12200000 | ATP5F1C, ITIH2, KIN |
| rs7067698 | 10 | 7795925 | 116858.4154 | 10p14 | chr10:6600000-12200000 | ATP5F1C, ITIH2, KIN |
| rs10905230 | 10 | 7821794 | 103894.6498 | 10p14 | chr10:6600000-12200000 | ATP5F1C, ITIH2, KIN |
| rs12770829 | 10 | 7842900 | 497359.5416 | 10p14 | chr10:6600000-12200000 | TAF3, ATP5C1, ITIH2, KIN, ATP5F1C |
| rs7091365 | 10 | 8078246 | 73317.78488 | 10p14 | chr10:6600000-12200000 | GATA3 |
| rs2503997 | 10 | 33215725 | 107922.2093 | 10p11.22 | chr10:31300000-34400000 | ITGB1, CCDC7 |
| rs78396600 | 10 | 118266459 | 107573.2085 | 10q25.3 | chr10:114900000-119100000 | PNLIP |
| rs3830026 | 10 | 118423572 | 104943.5434 | 10q25.3 | chr10:114900000-119100000 | PNLIPRP1 |
| rs6597996 | 11 | 674499 | 85068.73864 | 11p15.5 | chr11:0-2800000 | TMEM80, LMNTD2-AS1, LRRC56, DEAF1, HRAS, RNH1, IRF7, CDHR5, EPS8L2, PHRF1, SCT |
| rs7930974 | 11 | 122847880 | 119911.1298 | 11q24.1 | chr11:121200000-123900000 | BSX |
| rs11218912 | 11 | 122851155 | 133058.4895 | 11q24.1 | chr11:121200000-123900000 | BSX, JHY |
| rs1870761 | 11 | 122851563 | 93524.77054 | 11q24.1 | chr11:121200000-123900000 | BSX, JHY |
| rs77349614 | 12 | 51476387 | 292913.7214 | 12q13.12 | chr12:49100000-51500000 | CSRNP2 |
| rs11834113 | 12 | 93966218 | 736476.1239 | 12q22 | chr12:92600000-96200000 | SOCS2, SOCS2-AS1 |
| rs912784 | 13 | 46785521 | 149960.7832 | 13q14.13 | chr13:45800000-47300000 | LRRC63 |
| rs79530430 | 13 | 48652782 | 82715.56804 | 13q14.2 | chr13:47300000-50900000 | MED4 |
| rs41286118 | 13 | 76210890 | 133501.6461 | 13q22.2 | chr13:75400000-77200000 | UCHL3, LMO7-AS1 |
| rs34286592 | 16 | 29820480 | 109051.5463 | 16p11.2 | chr16:28100000-34600000 | INO80E, MAZ, PRRT2, NPIPB12, MVP, ASPHD1, KCTD13, DOC2A, CDIPTOSP, YPEL3, CDIPT, MAPK3, ZNF764, ZNF785 |
| rs749754 | 16 | 77469249 | 167531.6039 | 16q23.1 | chr16:74100000-79200000 | ADAMTS18 |
| rs13342232 | 17 | 6945940 | 3937431.481 | 17p13.1 | chr17:6500000-10700000 | RNASEK, SLC16A11 |
| rs11656323 | 17 | 7145117 | 577537.9843 | 17p13.1 | chr17:6500000-10700000 | GABARAP |
| rs222843 | 17 | 7145981 | 635663.1464 | 17p13.1 | chr17:6500000-10700000 | GABARAP |
| rs2074217 | 17 | 7150552 | 126800.6674 | 17p13.1 | chr17:6500000-10700000 | GABARAP |
| rs116903547 | 17 | 7162056 | 164876.8492 | 17p13.1 | chr17:6500000-10700000 | ENSG00000262302 |
| rs35776863 | 17 | 7226957 | 337426.6063 | 17p13.1 | chr17:6500000-10700000 | NEURL4 |
| rs2269762 | 17 | 7285831 | 165683.5814 | 17p13.1 | chr17:6500000-10700000 | TNK1 |
| rs17882626 | 17 | 7306727 | 670971.1067 | 17p13.1 | chr17:6500000-10700000 | TMEM256 |
| rs8069501 | 17 | 7394968 | 167818.0745 | 17p13.1 | chr17:6500000-10700000 | POLR2A |
| rs6608 | 17 | 7464413 | 93223.00225 | 17p13.1 | chr17:6500000-10700000 | SENP3 |
| rs11078697 | 17 | 7469229 | 344382.3351 | 17p13.1 | chr17:6500000-10700000 | SENP3 |
| rs28625968 | 17 | 7474539 | 152209.4066 | 17p13.1 | chr17:6500000-10700000 | EIF4A1 |
| rs9901673 | 17 | 7484101 | 94896.31986 | 17p13.1 | chr17:6500000-10700000 | CD68 |
| rs858520 | 17 | 7530271 | 127647.2784 | 17p13.1 | chr17:6500000-10700000 | SAT2 |
| rs6503227 | 17 | 9441453 | 102338.3526 | 17p13.1 | chr17:6500000-10700000 | STX8 |
| rs34654539 | 17 | 9525429 | 92989.25091 | 17p13.1 | chr17:6500000-10700000 | USP43 |
| rs9896200 | 17 | 9617718 | 108480.6657 | 17p13.1 | chr17:6500000-10700000 | USP43 |
| rs16958920 | 17 | 9795138 | 132016.8483 | 17p13.1 | chr17:6500000-10700000 | RCVRN |
| rs11079028 | 17 | 40126133 | 373318.9663 | 17q21.2 | chr17:38400000-40900000 | CNP |
| rs74647883 | 17 | 40153964 | 124290.9403 | 17q21.2 | chr17:38400000-40900000 | ZNF385C, NKIRAS2, C17orf113, CNP |
| rs12386058 | 17 | 40335086 | 93511.66617 | 17q21.2 | chr17:38400000-40900000 | MLX, KCNH4, EZH1, RETREG3 |
| rs1662336 | 18 | 3263391 | 104516.6566 | 18p11.31 | chr18:2900000-7100000 | MYL12B |
| rs7229123 | 18 | 3459378 | 151778.9927 | 18p11.31 | chr18:2900000-7100000 | TGIF1 |
| rs11085226 | 19 | 799770 | 119589.3058 | 19p13.3 | chr19:0-6900000 | PTBP1 |
| rs28384673 | 19 | 2271386 | 428549.2478 | 19p13.3 | chr19:0-6900000 | OAZ1 |
| rs281440 | 19 | 10400304 | 715635.5075 | 19p13.2 | chr19:6900000-13900000 | ICAM5 |
| rs17852402 | 19 | 10404519 | 120849.9671 | 19p13.2 | chr19:6900000-13900000 | ICAM5, ZGLP1 |
| rs75095909 | 19 | 10405734 | 143746.5908 | 19p13.2 | chr19:6900000-13900000 | ICAM3, ICAM5, FDX2, KEAP1 |
| rs13306513 | 19 | 11218226 | 109445.908 | 19p13.2 | chr19:6900000-13900000 | LDLR |
| rs5851 | 19 | 11832283 | 134740.5035 | 19p13.2 | chr19:6900000-13900000 | ACP5, ZNF823, ZNF763, ZNF833P, ZNF441 |
| rs117129463 | 19 | 11833719 | 641608.2471 | 19p13.2 | chr19:6900000-13900000 | ZNF823 |
| rs384148 | 19 | 11878095 | 303524.6353 | 19p13.2 | chr19:6900000-13900000 | ZNF69, ZNF440, ZNF491, ZNF439, ZNF441 |
| rs7624 | 19 | 35224470 | 535958.0779 | 19q13.11 | chr19:32400000-35500000 | SCGB2B2, SCGB1B2P, ZNF302, ZNF181 |
| rs7257305 | 19 | 35257192 | 93823.19053 | 19q13.11 | chr19:32400000-35500000 | ZNF302, ZNF599, SCGB1B2P, SCGB2B2 |
| rs17718432 | 19 | 35259048 | 122045.7758 | 19q13.11 | chr19:32400000-35500000 | ZNF181, ZNF599, SCGB2B2 |
| rs3746243 | 19 | 35449082 | 320314.4426 | 19q13.11 | chr19:32400000-35500000 | ZNF792 |
| rs2546027 | 19 | 35454215 | 277629.5011 | 19q13.11 | chr19:32400000-35500000 | GRAMD1A, SCN1B, ZNF792 |
| rs2073900 | 19 | 35741456 | 196884.348 | 19q13.12 | chr19:35500000-38300000 | LSR, FAM187B |
| rs67708190 | 19 | 36037148 | 594712.9265 | 19q13.12 | chr19:35500000-38300000 | TMEM147 |
| rs1165838 | 19 | 41075990 | 77533.9427 | 19q13.2 | chr19:38700000-43400000 | SHKBP1 |
| rs7248468 | 19 | 41240312 | 101073.4395 | 19q13.2 | chr19:38700000-43400000 | CYP2B7P, SNRPA, EGLN2, ITPKC, MIA, COQ8B, CYP2T1P, NUMBL |
| rs892596 | 19 | 45003267 | 97182.69254 | 19q13.31 | chr19:43400000-45200000 | ZNF234, ZNF180 |
| rs744193 | 20 | 21490151 | 90038.83385 | 20p11.22 | chr20:21300000-22300000 | NKX2-2 |
| rs80310989 | 20 | 43726528 | 437754.8719 | 20q13.12 | chr20:42100000-46400000 | KCNS1 |
| rs2531716 | 22 | 20004704 | 307569.5947 | 22q11.21 | chr22:17900000-22200000 | TANGO2 |

**S2 Table.**
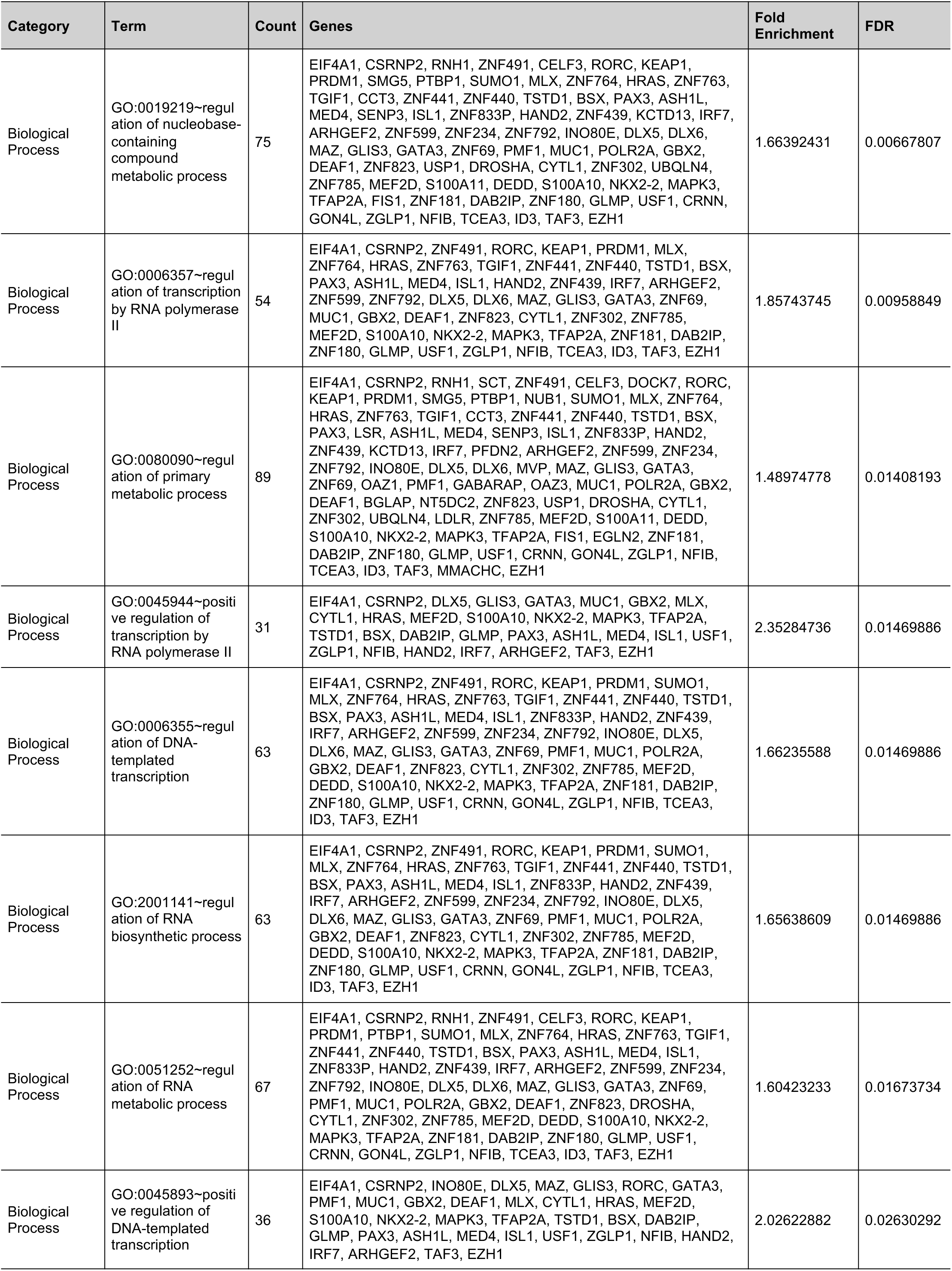
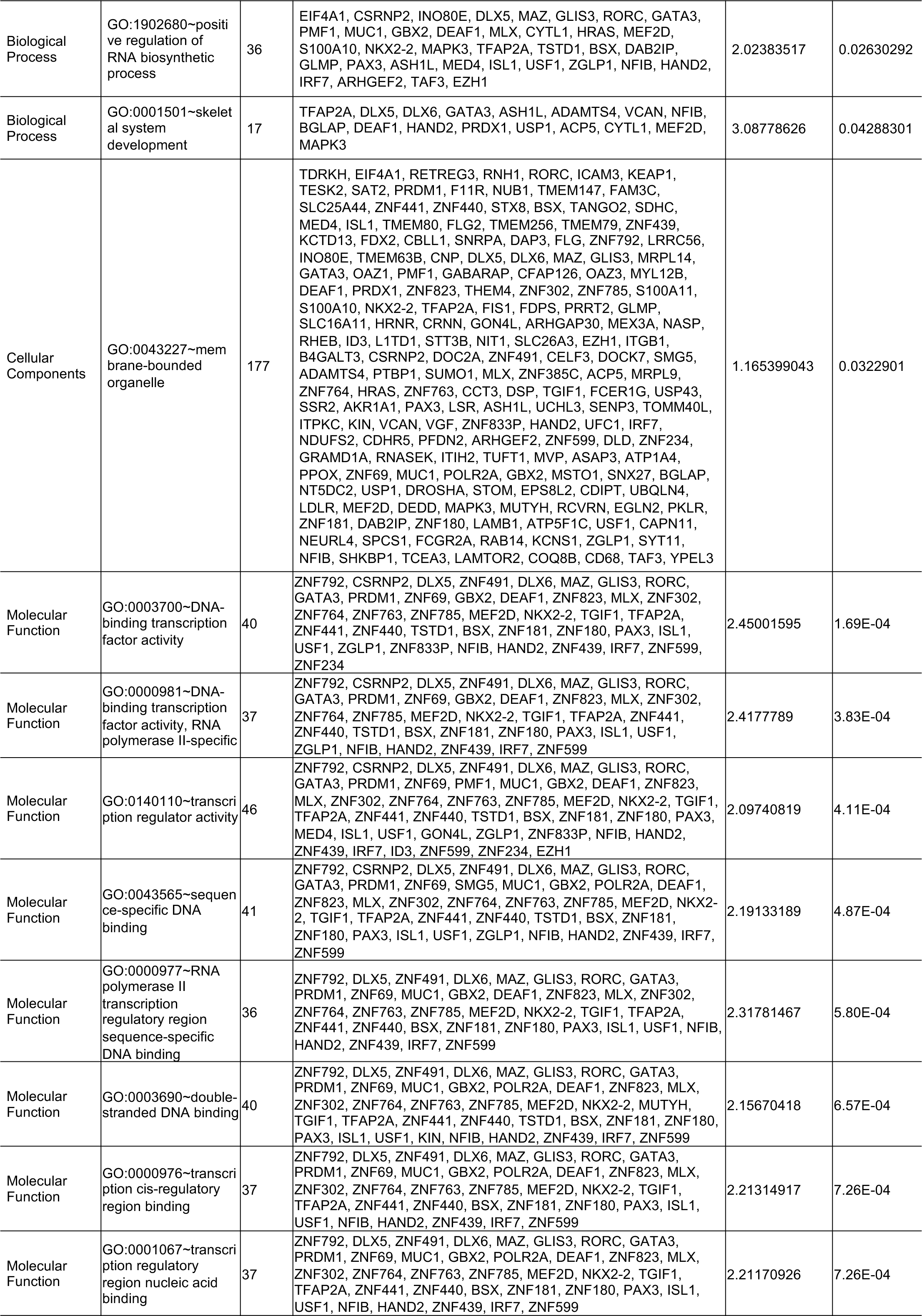
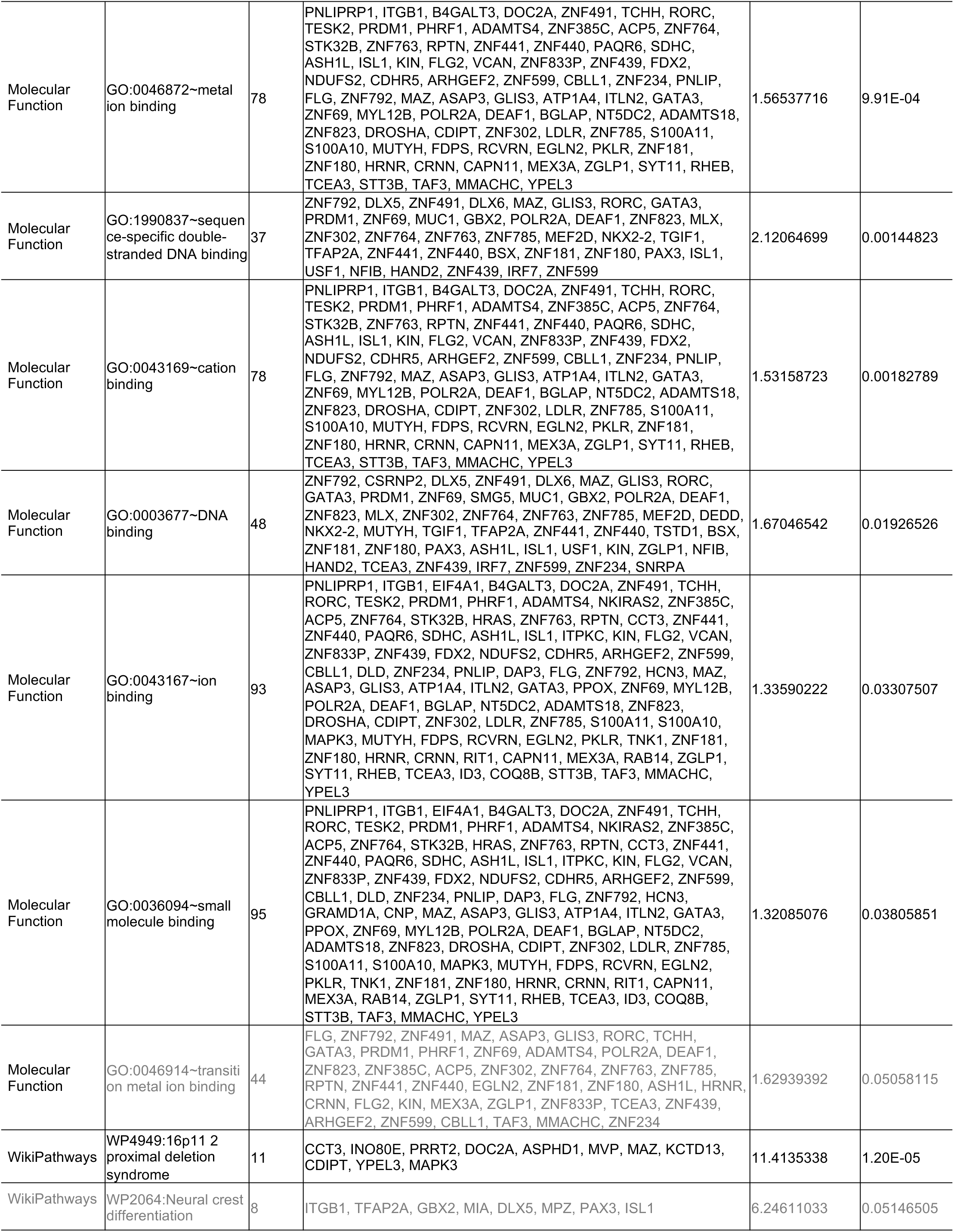
Enriched GO and Pathway terms. Enrichment analysis was performed on the 249 candidate genes using the DAVID online platform. Cutoff for enrichment was set at FDR = 0.05. Two marginally enriched terms (FDR = ∼0.05) are shown in grey text.. (D) Heatmap showing top marker genes of 11 annotated ectodermal clusters isolated from (B) and (C)

## Notes

### Competing Interest Statement

The authors have declared no competing interest.

